# MicroRNAs expressed from FSHR and aromatase genes target important ovarian functions

**DOI:** 10.1101/597054

**Authors:** Ilmatar Rooda, Birgitta Kaselt, Andres Salumets, Agne Velthut-Meikas

## Abstract

MicroRNAs (miRNAs) have known roles in the post-transcriptional regulation of various biological processes including ovarian follicle development. We have previously identified miRNAs from human pre-ovulatory granulosa cells that are expressed from the intronic regions of two key genes in normal follicular development: FSH receptor (FSHR) and CYP19A1, the latter encoding the aromatase enzyme. In the present study, we aim to identify the targets regulated by those two miRNAs: hsa-miR-548ba and hsa-miR-7973, respectively. The miRNAs of interest were endogenously expressed in KGN cell-line, gene expression changes were analyzed by Affymetrix microarray and confirmed by RT-qPCR. Potential miRNA-regulated sequences were further filtered from the obtained results by bioinformatic target prediction algorithms and validated for direct miRNA:mRNA binding by luciferase reporter assay. Our results verified Leukemia inhibitory factor receptor (LIFR), Phosphatase and tensin homolog (PTEN), Neogenin 1 (NEO1) and SP110 nuclear body protein (SP110) as target genes for hsa-miR-548ba. Hsa-miR-7973 target genes ADAM metallopeptidase domain 19 (ADAM19), Peroxidasin (PXDN) and Formin like 3 (FMNL3) also passed all verification steps. In conclusion we propose that hsa-miR-548ba may be involved in the regulation of follicle growth and activation via LIFR and PTEN. Hsa-miR-7973 may be implicated in the modulation of extracellular matrix and cell-cell interactions. Taken together, our results suggest that those two miRNAs of interest have important regulatory roles in granulosa cells and in follicle development in general.

**Summary sentence:** Confirmed targets of miRNAs hsa-miR-548ba and hsa-miR-7973 are involved in follicle recruitment, apoptosis, intercellular interactions and extracellular matrix remodeling pathways in KGN cells.

## Introduction

MicroRNAs (miRNAs) are short non-coding RNA molecules approximately 22 nucleotides in length [1]. MicroRNAs bind to their target mRNA 3’ untranslated region (3’UTR) at sites complementary to the miRNA 5’ seed region. MicroRNA biding to the target gene regulates gene expression by destabilizing the mRNA and/or inhibiting its translation [2]. In a few cases, an increase in gene expression has also been observed [3].

MicroRNAs have key role in the post-transcriptional regulation of various important biological processes, including cell proliferation, differentiation, apoptosis, and hormone biosynthesis and secretion [4]. Functional ovary requires precise coordination of follicle recruitment, selection and ovulation processes. Development of ovarian follicles is a complex process including oocyte maturation, granulosa cell proliferation and differentiation. MicroRNAs expressed in the ovary have regulative roles in ovarian follicle development and the expression of miRNAs in the ovary varies within the specific cell type and function [5]. The overall role of miRNAs in ovarian functions has been extensively studied in the mouse models. *Dicer1* conditional knock-out mice without the ability to process mature miRNAs in anti-Müllerian hormone expressing cells, including follicular granulosa cells, demonstrate abnormalities in ovulation, early embryonic development, and estrous cycles [6].

Granulosa cell functions are essential in follicular development, maturation, and atresia. Granulosa cells support oocyte growth via continuous bidirectional communication to ensure oocyte quality and developmental competence. Due to the close communication cumulus (CGC) and mural granulosa cells (MGC) may reflect the characteristics of the oocyte and an understanding of the patterns of expression and functions of miRNAs in those cells may lead to a better understanding of follicle maturation and its dysfunction [7,8]. Furthermore, granulosa cell miRNAs may serve as potential biological markers that could be used to increase the efficiency of assisted reproductive technologies by providing non-invasive means to assess oocyte quality and embryo survival potential [7].

MicroRNAs hsa-miR-548ba and hsa-miR-7973 were previously identified by deep sequencing of MGC and CGC populations isolated from women undergoing controlled ovarian stimulation and in vitro fertilization. Both miRNAs are of intronic origin: hsa-miR-548ba gene resides in the follicle stimulating hormone receptor (*FSHR*) gene and hsa-miR-7973 is located in the intron of the *CYP19A1* gene [9]. The regulatory mechanisms and target genes for those two miRNAs are currently not known.

Follicle stimulating hormone (FSH) activates time-related changes in granulosa cell gene expression by binding to FSHR promoting proliferation, differentiation, antrum formation, and oocyte maturation. Moreover, FSH stimulates aromatase expression and estrogens production [10,11]. Estrogens are produced by aromatization of androgens by aromatase enzyme encoded from *CYP19A1* gene [12]. Both FSHR and aromatase are crucial for follicle development and maturation [11].

The aim of the current study was to identify the target genes of hsa-miR-548ba and hsa-miR-7973 in human granulosa cells by using granulosa KGN cell line as a model [13]. The genomic locations of the miRNAs of interest in FSHR and aromatase genes give a hypothesis about important regulatory roles of these two miRNAs in granulosa cells regarding follicle development and function.

## Materials and methods

### Cell culture of KGN cell line

Granulosa tumor-like cell line KGN was used as a model for cell line experiments [13]. Cells were grown in L-glutamine-containing Dulbecco’s Modified Eagle’s medium with 10% fetal bovine serum, 100U/ml penicillin and 0.1mg/ml streptomycin on 60×15mm culture plates at 37°C and 5% CO_2_. Cells were harvested with 0.05% trypsin and 0.02% EDTA (all solutions from Naxo, Tartu, Estonia).

### Isolation of primary granulosa cells

The study was approved by the Ethics Committee of the University of Tartu and an informed consent was obtained from all participants. Granulosa cells were collected from eight women undergoing intra-cytoplasmic sperm injection (ICSI) due to male-factor infertility. Ovarian hormonal stimulation was conducted according to the GnRH antagonist (Cetrotide, Merck Serono, Geneva, Switzerland) protocol with the administration of recombinant FSH (Gonal-F, Merck Serono, or Puregon, Merck Sharp & Dohme Corp., Whitehouse Station, NJ, USA). All patients underwent ovarian puncture (OPU) of follicles ≥15 mm in size after 36 h of hCG administration (Ovitrelle, Merck Serono).

MGCs were obtained from follicular fluid after OPU following the manual removal of cumulus-oocyte complexes and CGC aggregates devoid of the oocyte. The fluid from all follicles of a patient was pooled, centrifuged at 450*g* for 10 minutes, followed by supernatant removal. The cells were separated on a 50% density gradient of PureSperm 100 (Nidacon; Mölndal, Sweden) in Universal IVF Medium (Origio; Jyllinge, Denmark), washed three times in Universal IVF Medium at 37°C, depleted of CD45-positive leukocytes according to the manufacturer’s suggested protocol (DynaMag and Dynabeads; Life Technologies, Carlsbad, California), lysed with QIAGEN miRNeasy Mini kit lysis buffer (QIAGEN, Hilden, Germany), and stored in liquid nitrogen for future use.

CGC were collected 4 h after OPU during oocyte denudation lasting up to 5 min with type IV-S hyaluronidase extracted from bovine testes (Sigma-Aldrich, St-Louis, MO, USA) and diluted in Sperm Preparation Medium (Origio). The CGC from all oocytes irrespective of their maturity were pooled and centrifuged at 450g for 5 min, the supernatant was discarded, and the cells were lysed and stored as described above.

### Cell line transfection

For microRNA transfection Lipofectamine RNAiMAX reagent (Invitrogen, Carlsbad, C and miRCURY LNA miRNA mimics hsa-miR-548ba, hsa-miR-7973 and control miRNA with no mammalian homologue cel-miR-39-3p [14,15] were used according to manufacturer’s protocol (Exiqon, Vedbaek, Denmark). MicroRNA mimics were transfected at previously optimized concentration of 12.5nM and 72 h time-point after transfection was tested.

For miRNA and luciferase vector co-transfection Lipofectamine 2000 reagent (Invitrogen) and previously specified miRCURY LNA miRNA mimics were used according to manufacturer’s protocol. MicroRNA final concentration in culture medium was 12.5nM or 50nM, while 100ng of vector was transfected.

### Cytotoxicity analysis

MicroRNA cytotoxicity analysis was performed using CytoTox-Glo Cytotoxicity Assay (Promega Corporation, Madison, Wisconsin, USA) according to the user manual. Five thousand cells were plated in each well of a white Greiner CELLSTAR 96-well plate (Sigma-Aldrich, St-Louis, MO, USA) and transfection was performed 24 h after plating. Non-transfected control samples were measured at 0, 24, 48, and 72 h. The cytotoxicity of cells transfected with either hsa-miR-548ba, hsa-miR-7973 or cel-miR-39-3p mimics was measured at 24, 48 and 72 h post-transfection. The above-mentioned assay uses a luminogenic peptide substrate to measure dead-cell protease activity, which is released from cells upon losing their membrane integrity. Luminescence was measured in two steps with Tecan GeniosPro luminometer (Tecan, Männedorf, Switzerland). Firstly, luminescence from dead cells and secondly luminescence from all cells after lysis was detected. Results were calculated as luminescence from all lysed cells minus luminescence from dead cells. All samples were measured in triplicates and the average value of three independent experiments was used in data analysis.

### RNA extraction

Cells were lysed in QIAzol lysis reagent (QIAGEN). For total RNA extraction miRNeasy Mini Kit (QIAGEN) was used. In addition, small fraction RNA (≤200 nucleotides) extraction was performed separately using RNeasy Mini Elute Cleanup Kit (QIAGEN). Both total and small RNA extraction was performed according to the user manual. RNA concentrations were measured using NanoDrop 2000c spectrophotometer (Thermo Fisher Scientific, Dreieich, Germany).

The quality of all RNA samples analyzed on Affymetrix microarray was evaluated using the Agilent 2100 Bioanalyzer system (Agilent Technologies, Waldbronn, Germany). All RNA samples were of high quality, with the RNA Integrity Number values between 9.2-9.8.

### Affymetrix microarray

Affymetrix microarray was performed at the Array and Analysis Facility at Uppsala Biomedical Center, Sweden. From each sample 250ng of total RNA were used to generate amplified and biotinylated sense-strand cDNA according to the GeneChip WT PLUS Reagent Kit User manual (Affymetrix, Santa Clara, CA). GeneChip Human Gene 2.0 ST Arrays were hybridized for 16h in 45°C incubator, rotated at 60rpm. Arrays were washed and stained using the Fluidics Station 450 and scanned using GeneChip® Scanner 3000 7G. Affymetrix microarray data is available at Gene Expression Omnibus repository, accession number GSE122731.

### Reverse transcription

For cDNA synthesis from mRNA SuperScript III First Strand Synthesis SuperMix (Invitrogen) was used according to user manual. 1μg of cell line and 500ng of primary granulosa cell RNA was used as input.

For microRNA detection cDNA was synthesized using miRCURY LNA™ Universal RT microRNA PCR Universal cDNA Synthesis Kit III (Exiqon). 200ng of small fraction RNA was used as input.

### Primer design

Primers used for RT-qPCR analysis were designed using NCBI primer-BLAST (http://www.ncbi.nlm.nih.gov/tools/primer-blast/). Preferred primers spanned exon-exon junctions and amplified all splice isoforms of the given gene. Primer sequences are presented in Supplementary Table IA.

Primers for amplifying the 3’UTR sequences of potential miRNA target genes were designed in Benchling (https://benchling.com/). Primer binding specificity was tested using NCBI primer-BLAST (Supplementary Table IB).

### RT-qPCR

The RT-qPCR analysis was carried out using SDS 2.3 software in 7900HT Fast Real-Time PCR System (Applied Biosystems, Foster City, California). For the detection of mRNA expression, Platinum SYBR Green qPCR SuperMix-UDG (Invitrogen) or Platinum SYBR Green qPCR SuperMix-UDG with ROX (Invitrogen) was used. cDNA from small RNA fraction was amplified by EXILENT SYBR Green Mastermix (Exiqon) according to the user manual. Each sample was run in triplicates on a 384-well plate. The specificity of amplified PCR products was determined by melt curve analysis.

### Luciferase reporter vector cloning and luciferase assay

The full length 3’UTR of hsa-miR-548ba potential target genes *BCL2L11, LIFR, NEO1, PTEN, RARB, SP110*, hsa-miR-7973 target genes *ADAM19, ATHL1, ATP6V1A, PXDN, FMNL3*, and the common target for both miRNAs *TGFBR2* were cloned into pmirGLO Dual-Luciferase vector (Promega) downstream of the Firefly luciferase gene. Target gene 3’UTR sequences were obtained from UCSC Genome Browser (https://genome.ucsc.edu/) with an exception of *FMNL3* where shorter isoform of 3’UTR was used. Bioinformatic prediction program TargetScan 7.1 [16] uses *FMNL3* 3’UTR isoform with length 7,874 nt, while in UCSC Genome Browser the length is 9,316 nt. Moreover, for *PTEN* also a shorter version of 3’UTR isoform was cloned which is used by miRDB bioinformatical target prediction program [17]. Potential target gene 3’UTR sequences and predicted miRNA binding sites are available in Supplementary Material. Primers for amplifying full length 3’UTR from genomic KGN cell line DNA and restriction enzymes used for cloning are presented in Supplementary Table IB.

Validation of hsa-miR-548ba and hsa-miR-7973 direct binding to target gene 3’UTR was performed by using Dual-Glo Luciferase assay system (Promega) according to the user manual. Luciferase signal was measured 24 h after miRNA and vector co-transfection. The relative luciferase activities were determined by calculating the ratio of Firefly luciferase activities over Renilla luciferase activities. All experiments were repeated three times as triplicates on a white Greiner CELLSTAR 96-well plate (Sigma-Aldrich). Cel-miR-39-3p was used as a negative control.

### Data analysis and statistics

#### a. Affymetrix microarray data analysis

Affymetrix microarray raw data was normalized in Expression Console, provided by Affymetrix (http://www.affymetrix.com), using the robust multi-array average (RMA) method. Differential gene expression analysis was carried out in the statistical computing language R (http://www.r-project.org) using packages available from the Bioconductor project (www.bioconductor.org) [18]. To detect differentially expressed genes between samples transfected with hsa-miR-548ba or hsa-miR-7973 and cel-miR-39-3p groups, an empirical Bayes moderated t-test was applied using the “limma” package [19]. The p-values were adjusted for multiple testing according to the method of Benjamini and Hochberg [20]. Statistical significance level was set at adjusted p < 0.01.

For cluster analysis of microarray data genes were ordered by the highest fold change expression difference upon hsa-miR-548ba transfection and Bioconductor package “heatmap” was used.

#### b. Gene ontology analysis

Analysis of over- and under-representation of Reactome pathways (version 58) was performed in the Panther classification system [21]. Genes differentially expressed (adjusted p < 0.01) on Affymetrix microarray upon transfecting KGN cells with either hsa-miR-548ba or hsa-miR-7973 mimics compared to the transfection with cel-miR-39-3p were used as input. Fisher’s exact test with Benjamini and Hochberg false-discovery rate (FDR) correction was used to test the statistical significance of the results and over- or under-represented pathways obtaining FDR < 0.05 are reported.

#### c. MicroRNA target gene prediction

For miRNA bioinformatic target prediction four web based programs were used (DIANA microT v 3.0 [22], microT CDS v5.0 [23], TargetScan 7.1 [16] and miRDB [17]). Gene was considered as a potential miRNA target in case it was predicted by at least two programs out of four and its gene expression fold change according to microarray analysis was ≥ log2(|0.3|).

#### d. RT-qPCR

The gene-specific mRNA expression Ct values were normalized against *GAPDH* expression and miRNA expression levels were normalized for internal control hsa-miR-132-3p using the 2^-ΔΔCt^ method of relative quantification [24]. Experiments were run in technical triplicates three times. Results are shown as average ± SD (standard deviation). Statistical significance was calculated by two-tailed Student t-test in Microsoft Office Excel 2017. Statistical significance level was set at p < 0.05.

#### e. Luciferase assay

The relative luciferase activities were determined by calculating the ratio of Firefly luciferase relative light units (RLU) over Renilla luciferase RLU. All experiments were run independently three times as triplicates. Results are shown as average ± SE (standard error). Statistical significance was calculated by one-tailed Student t-test in Microsoft Office Excel 2017. Statistical significance level was set at p < 0.05.

## Results

### Global gene expression changes upon transient expression of hsa-miR-548ba and hsa-miR-7973 in KGN cells

The first aim of the current study was to evaluate the effect of miRNA transfection on the global gene expression change in human granulosa cell line KGN. In non-transfected KGN cells the expression levels of hsa-miR-548ba and hsa-miR-7973 barely reached the detection limit (Supplementary Figure 1). After optimization experiments (data not shown), the transfection of 12.5nM miRNA mimic lead to considerably higher expression levels in comparison to primary granulosa cells (Supplementary Figure 1). However, such level of over-expression did not influence cell viability or proliferation rate (Supplementary Figure 2).

Genome-wide gene expression changes upon miRNA transfection were studied on Affymetrix GeneChip Human Gene 2.0 ST Array. The results demonstrated that upon hsa-miR-548ba transfection the expression level of 1,474 and upon hsa-miR-7973 the expression level of 1,552 genes changed with statistical significance (adjusted p-value <0.01, Supplementary Table IIA and IIC). From those genes 1,015 were regulated by both miRNAs, 459 genes only by hsa-miR-548ba and 537 by hsa-miR-7973. Gene expression changes were calculated as compared to the control samples transfected with miRNA cel-miR-39-3p that presumably has no target sequences in human cells.

Cluster analysis of microarray results revealed that control samples expressing cel-miR-39-3p grouped separately from samples transfected with hsa-miR-548ba and hsa-miR-7973 (Figure 1). Cells expressing the human miRNAs under study had more similar gene expression patterns compared to samples transfected with cel-miR-39-3p. This is also confirmed by the overlapping number of commonly regulated genes.

**Figure 1.**
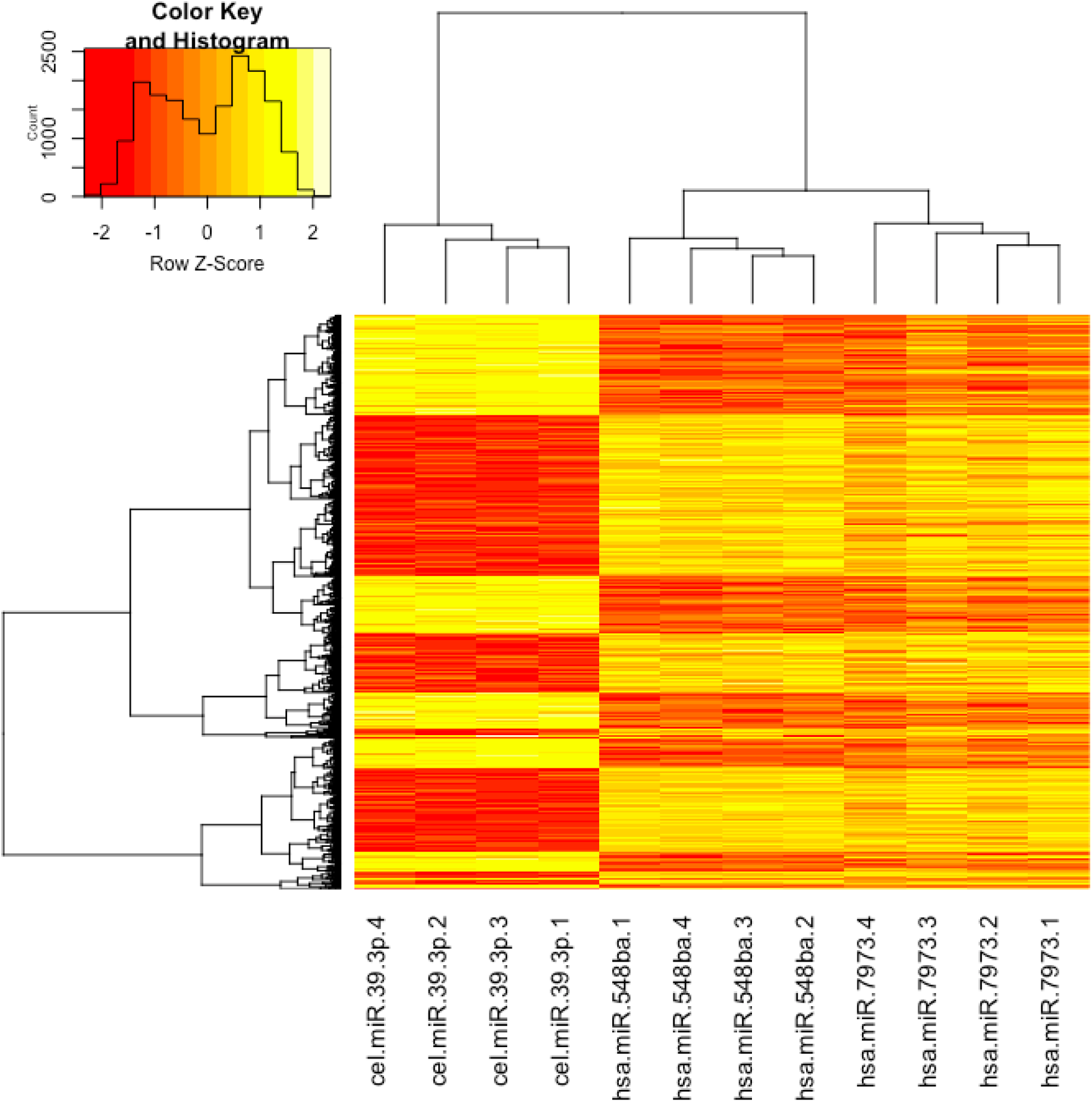
Cluster analysis of gene expression changes upon transfection of KGN cells with cel-miR-39p, hsa-miR-548ba or hsa-miR-7973 miRNA mimic. Gene expression changes were analyzed 72 h after transfection on Affymetrix microarray. Only statistically significant results are presented (adjusted p-value < 0.01).

Transfection of KGN cells with hsa-miR-548ba and hsa-miR-7973 leads to the regulation of several common as well as unique signaling pathways (Supplementary Table IIIA and IIIB). Genes differentially expressed upon hsa-miR-548ba transfection are over-represented in syndecan interactions, glycosaminoglycan and fatty acid metabolism, unfolded protein response, G protein-coupled receptor (GPCR) and ephrin signaling among others (Supplementary Table IIIA). Expression of hsa-miR-7973 in KGN cells resulted in the regulation of genes involved in signaling by netrin-1, EGFR, PDGF, and TGF-beta receptor complex, as well as pathways related to the immune system (Supplementary Table IIIB). As several genes were regulated by both miRNAs, common targeted pathways include cholesterol biosynthesis and amino acid transport across cell membrane (Supplementary Table IIIA and IIIB).

### Bioinformatic target prediction

Microarray analysis results may partly demonstrate secondary effects on mRNA expression: gene expression changes triggered by primary miRNA targets. Therefore, to discriminate between primary and secondary target genes the differential expression results obtained from the microarray analysis method were compared to potential miRNA target mRNA-s predicted by bioinformatic algorithms. However, bioinformatic prediction may also provide false positive hits. In order to minimize such false positive findings, four target prediction programs were used: DIANA microT v 3.0 [22], microT CDS v5.0 [23], TargetScan 7.1 [16] and miRDB [17]. A gene was considered as a potential true target for a miRNA when it was positively predicted by at least two programs out of four.

The overlap of statistically significant microarray results and bioinformatically predicted target genes are presented in Supplementary Table IIB and IID. Shortly, hsa-miR-548ba and hsa-miR-7973 potentially target the mRNA-s of 76 and 58 genes, respectively. Both miRNAs also share one common target mRNA of *TGFBR2* gene.

### Validation of microarray results by RT-qPCR

Sixteen potential target genes from the overlapping list of microarray and bioinformatic target prediction results were validated by RT-qPCR: 8 potential targets of hsa-miR-548ba and 7 of hsa-miR-7973 and their common target *TGFBR2*. The list of validated genes was selected according to published data linking the molecular functions of these proteins to their potential importance in folliculogenesis (Table I). As a result, *BCL2L11, LIFR, NEO1, PTEN* and *SP110* were statistically significantly down-regulated at mRNA level upon cell transfection with hsa-miR-548ba mimic (Figure 2A). *RARB* was down-regulated with borderline statistical significance (p=0.057, Figure 2A). From the list of potential hsa-miR-7973 targets the expression levels of *ADAM19, ATHL1, ATP6V1A, FMNL3* and *PXDN* were statistically significantly decreased (Figure 2B). The expression change of the common target gene *TGFBR2* was confirmed only in case of hsa-miR-7973 transient expression (Figure 2B).

**Figure 2.**
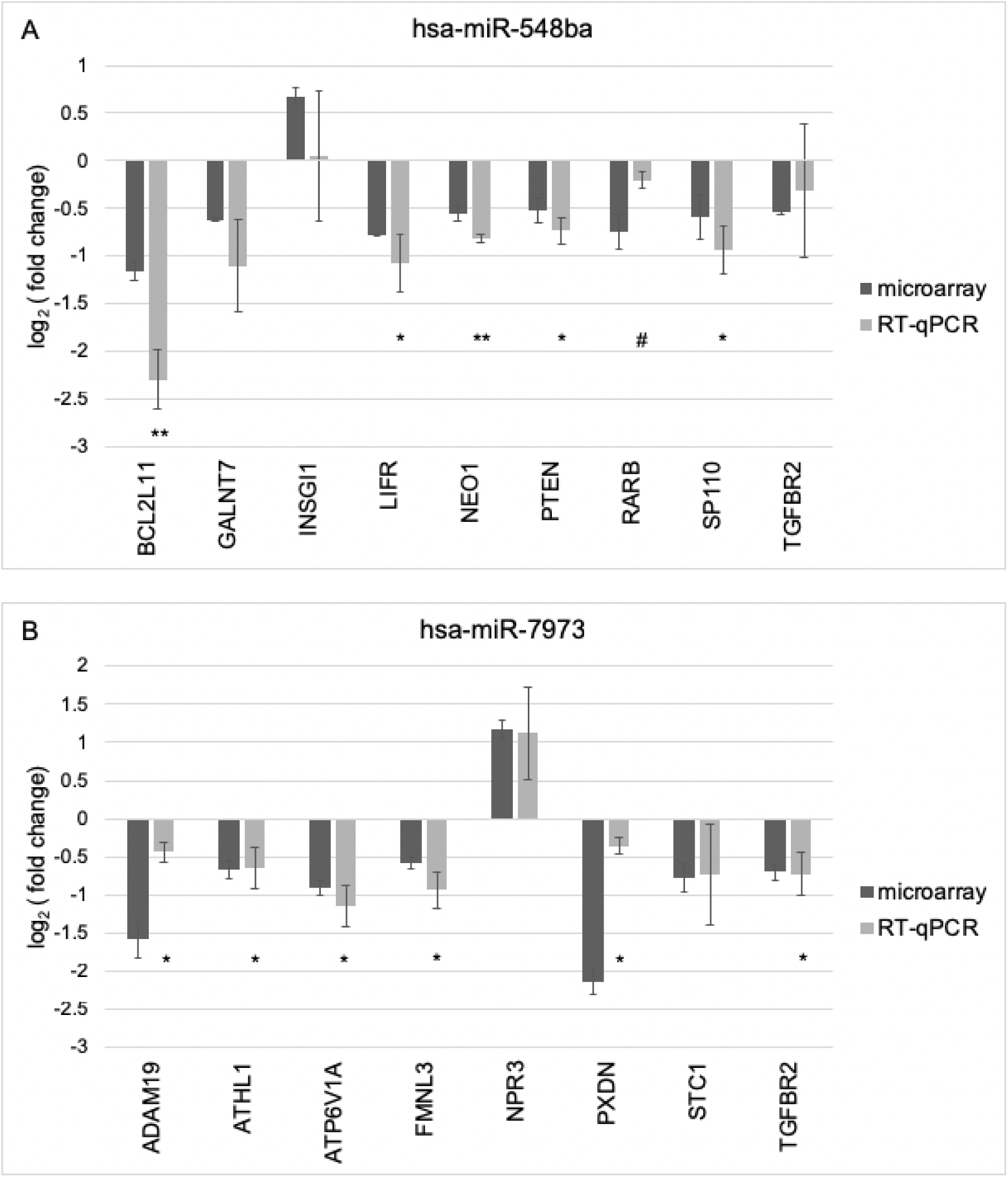
Validation of microarray results by RT-qPCR. A) Potential target genes of hsa-miR-548ba and B) Potential target genes of hsa-miR-7973. Gene expression change is calculated by comparing expression levels to samples transfected with control miRNA cel-miR-39-3p. Results are displayed as average fold change ±SD on log2 scale, (^#^p = 0.057, *p < 0.05; **p < 0.01; Student t-test). All presented microarray results were statistically significant (adjusted p < 0.01).

**Table 1.**
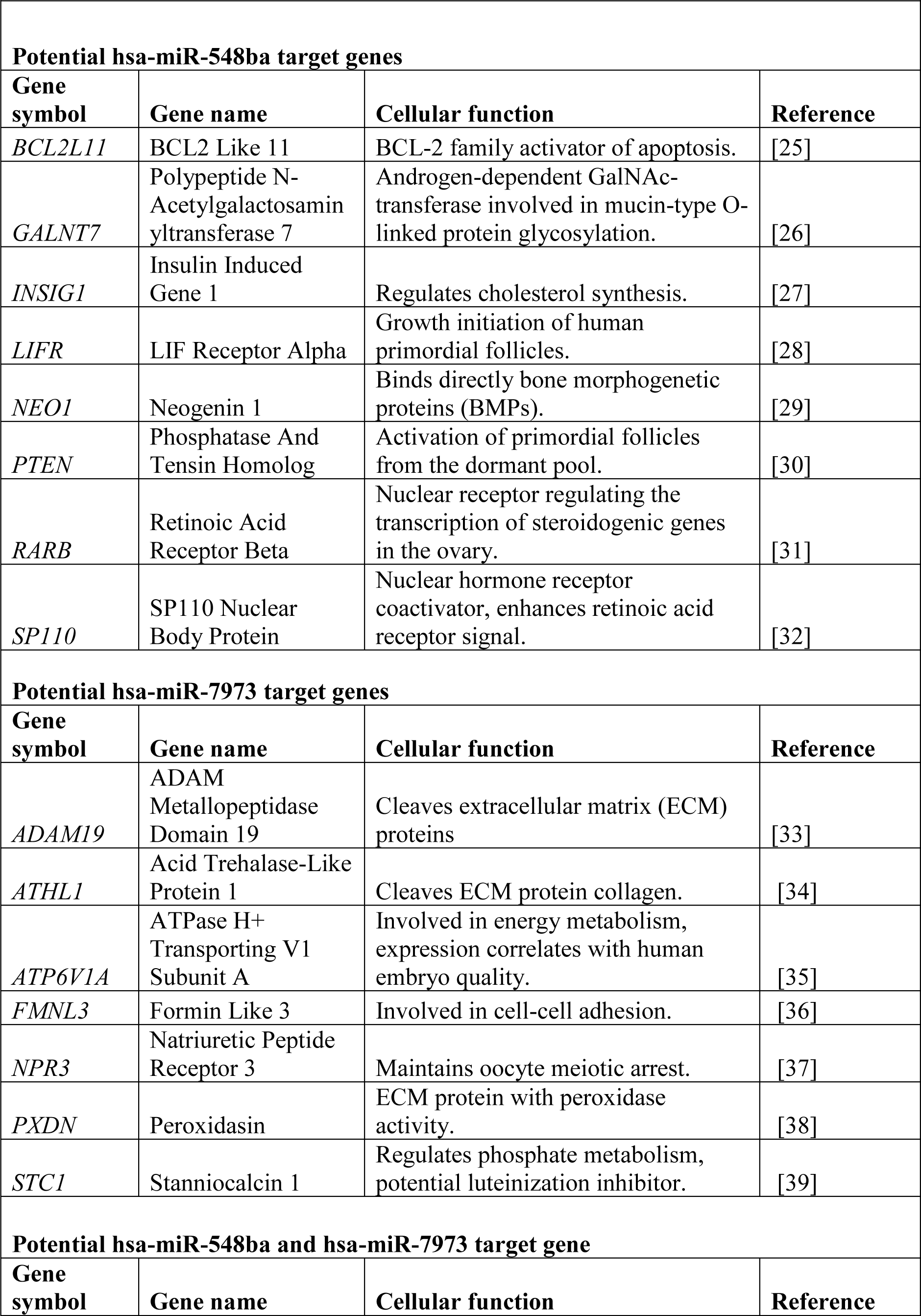
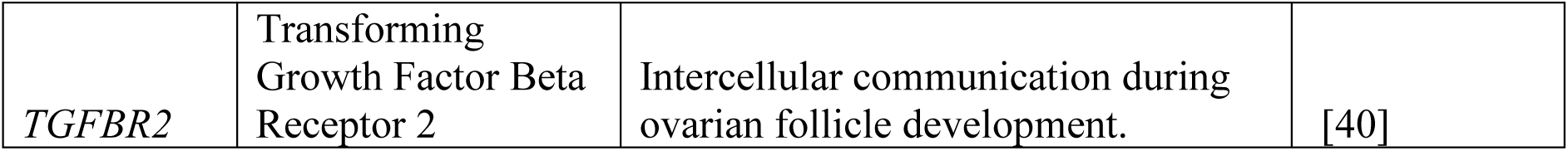
Potential target genes of hsa-miR-548ba and hsa-miR-7973 used in validation studies and their cellular functions.

### MicroRNA target binding validation by luciferase reporter assay

Potential miRNA targets that demonstrated significant gene expression change by microarray (p < 0.01) and RT-qPCR methods (p < 0.05) were further validated for direct miRNA:mRNA binding by luciferase reporter assay. Shortly, hsa-miR-548ba potential target genes: *BCL2L11, LIFR, NEO1, PTEN, RARB* and *SP110*, and hsa-miR-7973 target genes: *ADAM19, ATHL1, ATP6V1A, FMNL3* and *PXDN*. Although *TGFBR2* gene expression change was not confirmed by RT-qPCR in hsa-miR-548ba transfected cells, binding of its 3’UTR to both miRNAs under study was nevertheless assessed.

Potential target gene *PTEN* was tested with two versions of 3’UTR lengths: 3’UTR length obtained from UCSC Genome browser and second from miRDB bioinformatical prediction program, marked as *PTEN* long and *PTEN* short, respectively. pmirGLO-3’UTR-PTEN vector and hsa-miR-21-5p were used as a positive control for miRNA:mRNA binding to confirm our luciferase assay reliability as *PTEN* has been previously shown to be a target for hsa-miR-21-5p [41]. Upon direct binding of miRNA to the 3’UTR sequence of its target mRNA, reduction in the measured luciferase signal is expected. Such suppression of luciferase signal upon hsa-miR-21-5p binding to *PTEN* 3’UTR is demonstrated in Supplementary Figure 3 for both long and short variants.

Luciferase assay results confirmed the direct binding of hsa-miR-548ba to *LIFR, NEO1, PTEN* and *SP110* 3’UTR sequences. Hsa-miR-548ba bound to its potential target *PTEN* 3’UTR sequence only when the longer version was used (Figure 3A). From the tested potential targets of hsa-miR-7973 direct binding occurred on the 3’UTR of *ADAM19, FMNL3* and *PXDN* (Figure 3B).

**Figure 3.**
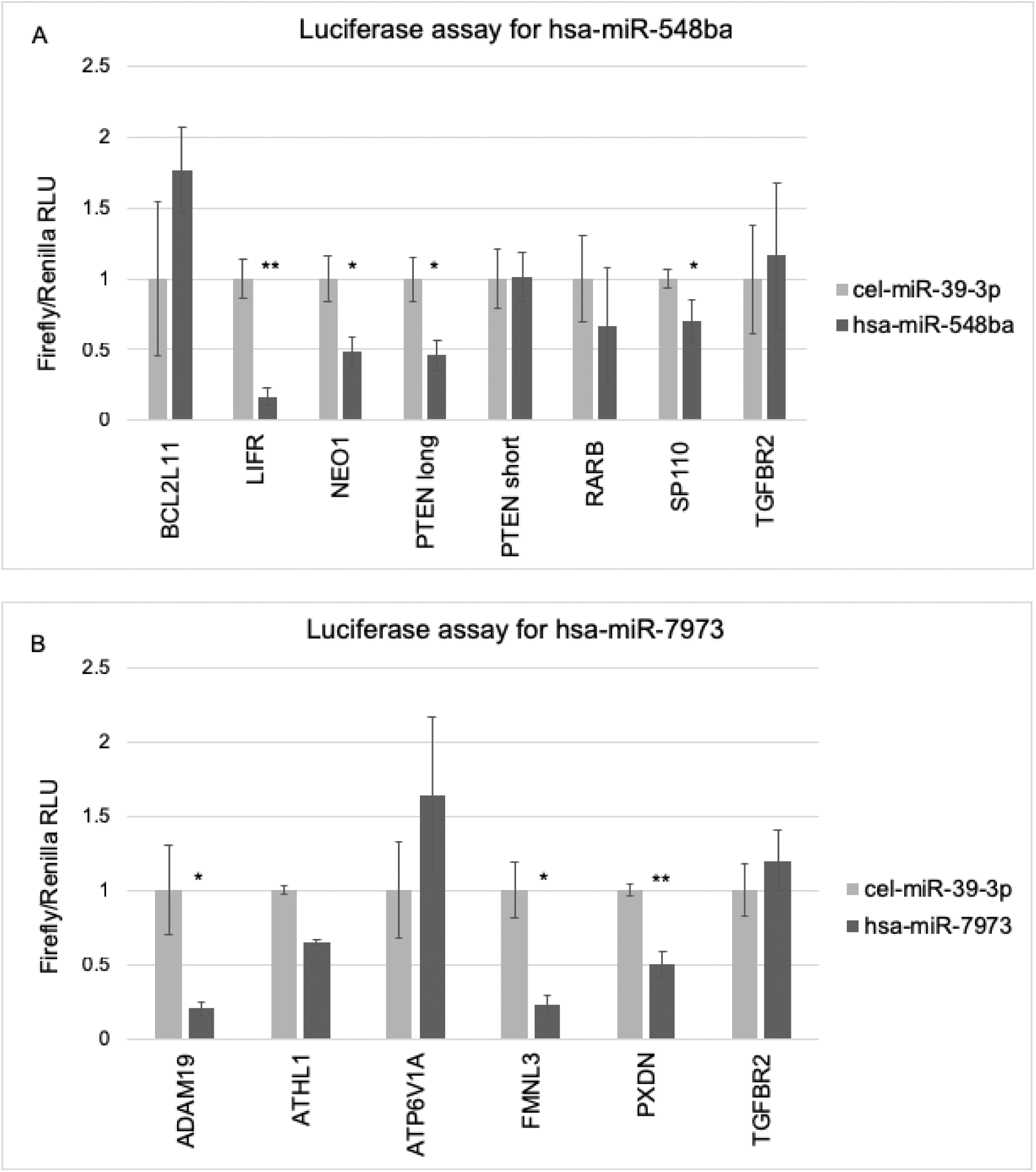
Validation of miRNA binding on the 3’UTR sequences of their potential target mRNAs by luciferase reporter assay. A) hsa-miR-548ba potential target genes. B) hsa-miR-7973 potential target genes. Results are shown as average normalized luciferase signal ±SEM, (*p < 0.05; **p < 0.01; Student one-tailed t-test). RLU – relative light unit.

## Discussion

Here we present the first functional analysis of miRNAs hsa-miR-548ba and hsa-miR-7973 by identifying their target genes in KGN cells using Affymetrix microarray, RT-qPCR and luciferase assay.

The miRNAs of interest: hsa-miR-548ba and hsa-miR-7973, were both identified for the first time from human ovaries by deep sequencing approach of primary granulosa cells [9]. In the described functional experiments we used granulosa tumor-like KGN cell line as granulosa cell model that has been previously used to understand the regulation of steroidogenesis, cell growth and apoptosis in human granulosa cells [13]. KGN cells express functional FSH receptor and high level of aromatase, encoded from miRNA hsa-miR-548ba and hsa-miR-7973 host genes *FSHR* and *CYP19A1*, respectively. KGN cell line has also been previously used for miRNA target identification and luciferase assay studies for direct miRNA:mRNA interactions [42].

MicroRNAs are estimated to regulate 60% of all protein-coding genes and a single miRNA may regulate up to 400 different target mRNAs [43]. Our microarray results demonstrated 1,474 differentially expressed genes upon hsa-miR-548ba transfection, and 1,552 genes potentially regulated by hsa-miR-7973. From differentially expressed genes 1,015 were regulated by both miRNAs, 459 genes only by hsa-miR-548ba and 537 by hsa-miR-7973. Gene ontology analysis from hsa-miR-548ba regulated genes showed enrichment in fatty acid metabolism and ephrin signaling pathways and over-expression of hsa-miR-7973 showed enrichment in TGF beta receptor complex and immune system pathways. Commonly regulated genes by these two miRNAs targeted cholesterol biosynthesis pathway. Cholesterol is a major component for steroidogenesis and sources of cholesterol for ovarian granulosa cells include plasma lipoproteins and de novo-synthesis [44]. Therefore, the regulation of cholesterol biosynthesis pathway by the miRNAs under study may influence steroid hormone production, contributing to the follicular developmental stage-dependent estradiol and progesterone production.

Although, miRNAs may regulate several hundreds of target genes, microarray results do not differentiate primary targets from secondary regulatory effects: i.e. the genes regulated by primary miRNA targets. Such secondary regulatory effects are common in over-expression conditions and large data-sets obtained by genome-wide methods like microarray gene expression analysis [45]. The large number of commonly regulated genes by both miRNAs analyzed in the present study can be also explained by secondary effects on gene expression, as miRNA transfection and over-expression can potentially lead to the saturation of RNA silencing complex (RISC) which results with altered gene expression regulation [46]. Moreover, miRNAs may target transcription factors influencing the expression of a number of secondary target genes [45]. To better distinguish primary miRNA targets from secondary ones, microarray results were compared to bioinformatically predicted target genes.

MicroRNAs are known to have regulatory roles in many stages of follicle development and granulosa cell functions [5]. To investigate the role of the two miRNAs of interest, the overlapping part of microarray and bioinformatically predicted potential targets were filtered for genes with a known role in ovarian function and tested for miRNA:mRNA binding effects on protein expression.

For direct miRNA:mRNA binding and protein expression studies luciferase assay reporter vectors were cloned with potential target gene 3’UTR sequences. In this study full length 3’UTR sequences were used, except for *PTEN* in case of which bioinformatic prediction algorithms do not use consistent 3’UTR sequence lengths and thus two different sequences were used. We chose to test full-length sequences for most of the target genes as we lacked the knowledge of functional miRNA target regions and the exact biding site identification within each potential target gene was not the goal of the current study. Furthermore, the secondary structure of 3’UTR could also determine the accessibility of miRNA binding leading to our choice of using full length 3’UTRs. Testing a fragment of 3’UTR may easily lead to false results due to the different secondary structure from the full length sequence [47]. Furthermore, 3’UTR central regions are more likely to be incorporated to hairpin structures which could lead to concealing the target sites of miRNAs. This is also supported by different binding site efficiencies: miRNA sites near the ends of the 3’UTR are more effective than sites in the center of the 3’UTR [47,48]. Direct binding assay confirmed *LIFR, PTEN, NEO1* and *SP110* as hsa-miR-548ba targets. For target gene *PTEN* direct binding occurred only with the longer 3’UTR sequence, although bioinformatic algorithms used in this work predicted target sequences into the shorter 3’UTR isoform. Such results can be explained by the differences in the secondary structure mentioned above. Moreover, bioinformatical programs may not predict all the potential target sequences for miRNA hsa-miR-548ba. Taken together, target gene 3’UTR length is critical for miRNA binding to its target sequences in direct binding assays.

LIFR is a receptor for leukemia inhibitory factor (LIF). LIF has been shown to promote primordial to primary follicle transition in the rat ovary [49]. In cattle, follicle atresia is regulated by LIF-STAT3 pathway which can be reversed with FSH administration [50]. In humans, the role of LIF and its receptor in the ovary is not well characterized, but LIF and LIFR mRNA expression in human ovarian samples are consistent with the concept that LIF might be involved in growth initiation of human primordial follicles through its receptor [28].

Second direct target gene for hsa-miR-548ba, *PTEN*, is involved in primordial follicle dormancy and activation. The recruitment of primordial follicles is regulated by highly controlled mechanisms and the critical role held by PI3K-Akt signaling pathway. This pathway is negatively regulated by PTEN and it is showed that in mice lacking *PTEN* the entire primordial follicle pool becomes activated [30]. In bovine the levels of PI3K-Akt signaling pathway components in MGC and CGC have a correlation with oocyte competence. Decreased PTEN expression and increased expression of PTEN downregulating miRNAs lead to the activation of PI3K-Akt pathway and correlates with the number of high-quality oocytes [51]. Therefore, the timing of PTEN expression regulation is important for follicle development and oocyte competence. Although hsa-miR-548ba expression pattern throughout the follicle development is currently unknown, a similar regulation of PTEN by hsa-miR-548ba may have an important role in the determination of oocyte developmental competence.

NEO1 is a member of immunoglobulin superfamily and is a receptor for repulsive guidance molecules, netrins, and is involved in bone morphogenetic protein (BMP) signaling pathways by directly binding BMPs [29]. Moreover, NEO1 activates RAC1-PI3K-AKT signaling pathway in human gastric cancer cells [52]. In mouse ovary, Rac1 is involved in primordial follicle formation by inducing nuclear transport of SMAD3 [53]. As mentioned above, PI3K-AKT signaling pathway have crucial roles to balance follicle growth suppression, activation and progression also in humans [54] and BMPs have important role in follicle development including primordial germ cell development, oocyte-somatic cell interactions and modulating of COC formation and expansion [55].

The role of SP110 in the ovary is less clear. It has been suggested that SP110 may function as a nuclear hormone receptor transcriptional coactivator via binding to the retinoic acid receptor alpha (RARA) response element [32]. Retinoids are essential for steroid production and it has been demonstrated that bovine CGCs express active RARA [56]. The role of SP110 in human granulosa cells has to be further investigated, but SP110 regulation by miRNA hsa-miR-548ba may influence retinoid signalization.

Moreover, direct hsa-miR-548ba targets *PTEN, NEO1* and potential secondary target *BCL2L11* are also involved in regulating apoptosis [25,57,58]. However, the role of hsa-miR-548ba in modulating apoptotic pathways in the human ovary needs to be investigated further. In the current study, exogenous expression of hsa-miR-548ba did not lead to changes in overall cellular viability during 72 h. Studies involving long-term ovarian culture conditions could shed more light on the importance of hsa-miR-548ba on cell viability.

The confirmed targets of hsa-miR-7973 according to our study were *ADAM19, PXDN* and *FMNL3*, all of which have been shown to be involved in ECM modulation and cell-cell interactions [33,38]. ADAM19 cleaves extracellular matrix (ECM) proteins, growth factors and cytokines like neuregulin and plays an essential role in embryo implantation by remodeling ECM [33]. Neuregulin 1 (NRG1) is expressed in granulosa cells and induced by luteinizing hormone (LH). NRG1 regulates the meiotic resumption time of the oocyte leading to proper oocyte maturation and developmental competence in cultured COCs [59]. The down-regulation of ADAM19 expression by miRNA hsa-miR-7973 could lead to lower cleavage rate of NRG1. Therefore, it is important to know the timing of miRNA hsa-miR-7973 regulation of ADAM19 and how this influences NRG1 cleavage levels.

PXDN is a peroxidase previously investigated in the context of ovarian cancer where the knockdown of this gene inhibits cellular proliferation, invasion and migration of ovarian cancer cells [38]. Moreover, PXDN may promote proliferation of ovarian cancer cells by regulating the activity of PI3K-AKT signaling pathway [38]. The role of PXDN in follicular cells still needs further clarification.

Regulation of FMNL3 is not previously investigated in the context of ovary but this protein has been shown to be involved in cytoskeletal organization, cell morphology, migration and cell-cell adhesion [36,60].

Genes with negative results from luciferase assay can be considered as secondary targets of miRNAs of interest but we have to keep in mind that the experiments were performed with reporter vectors and in cell line conditions that may not fully reflect physiological conditions in primary cells. Negative binding results may be the outcome of multiple biological reasons. For example, miRNA target recognition may require multiple transacting factors which leads to targeting only in specific cellular contexts. Also the alternative polyadenylation of 3’UTRs in reporter vectors may increase the false-negative results [61]. In addition, many possible target genes were excluded in this study due to the negative result from microarray gene expression analysis using the cell line model and were thus not tested in the direct binding assay. Therefore, in another experimental setting those genes may still be proven as direct targets, leading to the conclusion that the list of validated target genes for these two miRNAs of interest is not final.

Overall, we confirmed several targets for miRNA hsa-miR-548ba and hsa-miR-7973. According to the previously known functions of their target genes we suggest that those two miRNAs may have potential regulatory roles in granulosa cell gene expression and in follicle development in general.

## Supporting information

Supplementary Table III

## Acknowledgments

We are thankful to professor Markku Heikinheimo from the Institute of Clinical Medicine, University of Helsinki for providing KGN cell-line for this project. We thank Hanna Göransson Kultima, from the Array and Analysis Facility, SciLifeLab Uppsala, Sweden for her assistance in microarray data analysis.

## Supplemental Data Legends

**Supplementary Figure 1.**
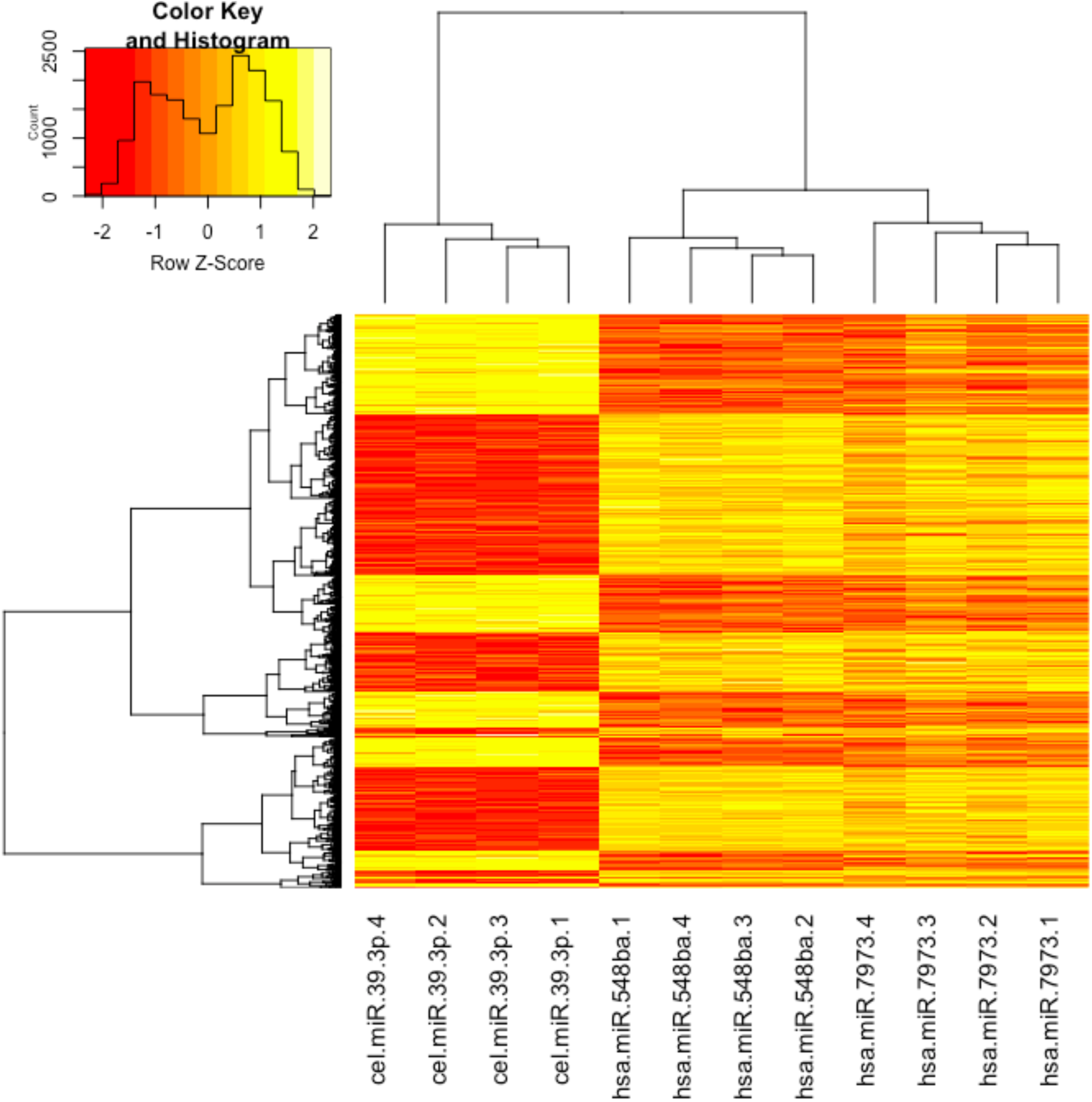
Transient hsa-miR-548ba and hsa-miR-7973 expression in KGN cell-line compared to primary human granulosa cells. For positive control human mural (MGC) and cumulus granulosa cell (CGC) RNA was used (n=8). miRNA expression in KGN cells is shown as average expression of four parallel samples ± SD on log2 scale. miRNA expression levels were normalized against miRNA hsa-miR-132-3p levels. Transfected KGN samples are indicated with + sign in the legend (yellow and pink color).

**Supplementary Figure 2.**
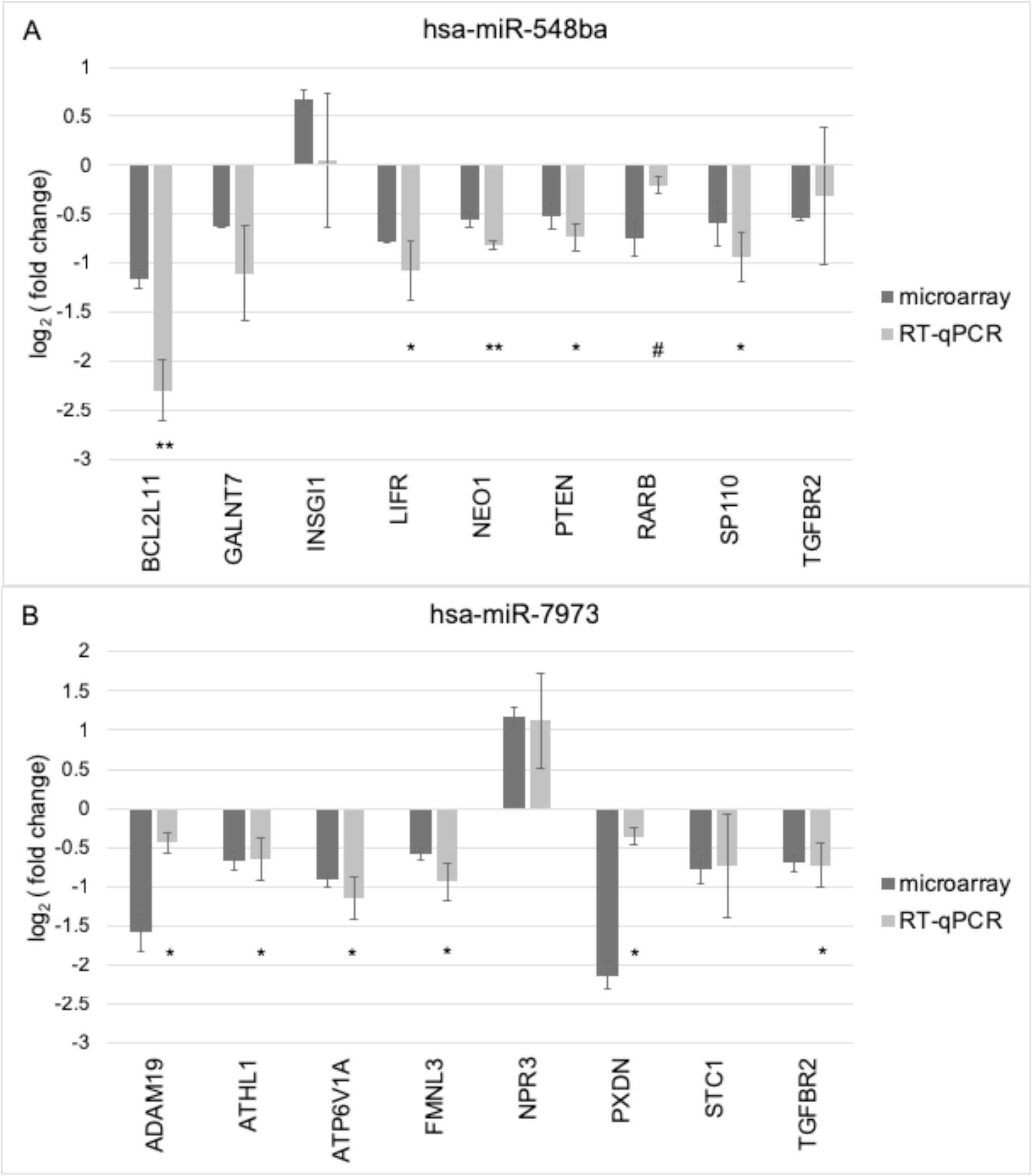
KGN cell viability upon transient miRNA expression. Luminescence signal was measured from cells transfected with hsa-miR-548ba, hsa-miR-7973 or control miRNA cel-miR-39-3p at three time-points: 24, 48 and 72 hours. Signal from non-transfected cells (control) were measured at 0, 24, 48 and 72 hours. The indicated value refers to normalized luminescence: signal from all lysed cells minus signal from dead cells. Results are shown as average of 3 independent experiment with ± SD. Student t-test p-values were above 0.05, no significance difference in living cell number were detected between non-transfected control samples and transfected cells.

**Supplementary Figure 3.**
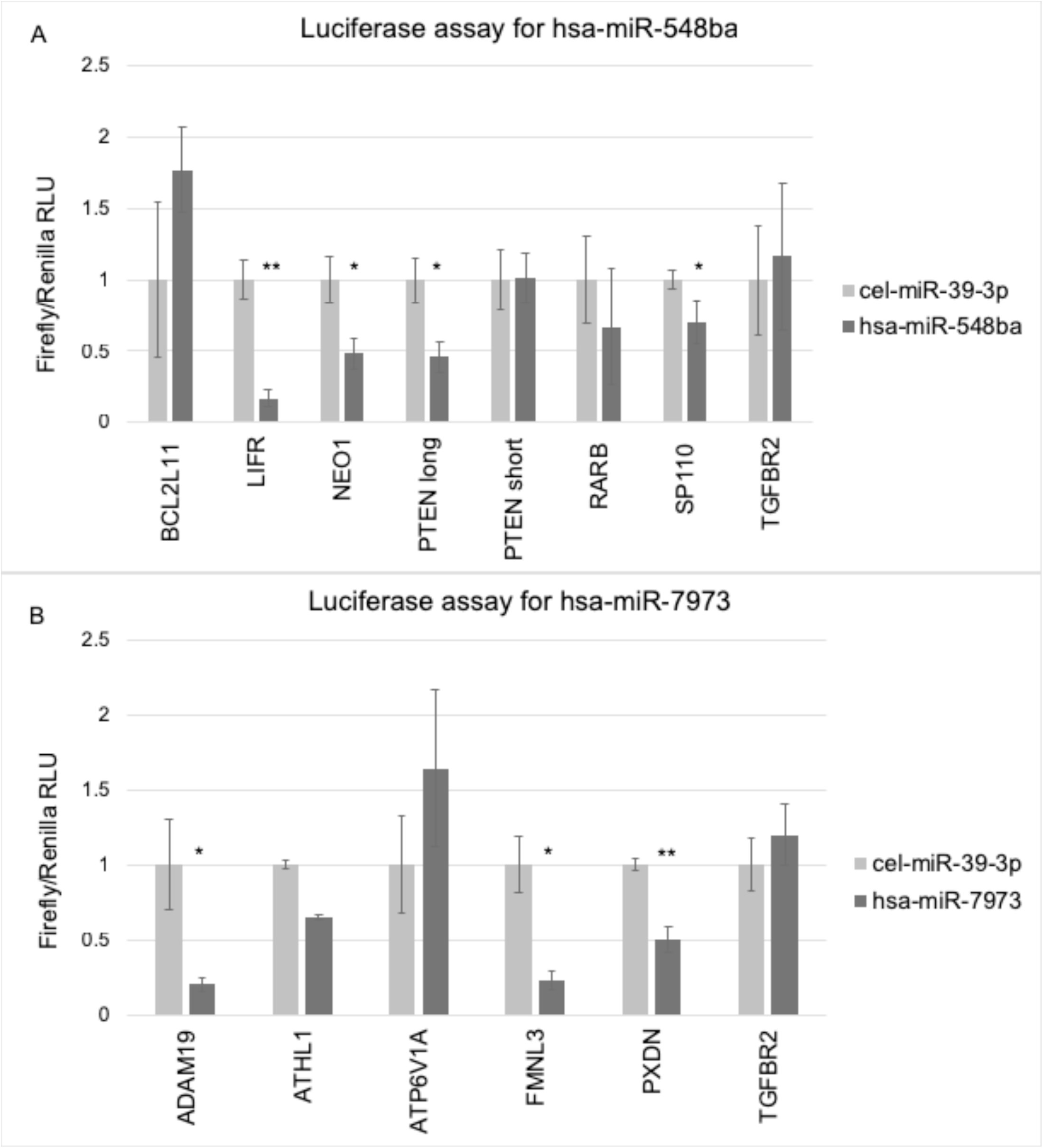
Suppression of luciferase signal upon hsa-miR-21-5p binding to PTEN 3’UTR. pmirGLO-3’UTR-PTEN vector and hsa-miR-21-5p were used as a positive control for miRNA:mRNA binding to confirm the luciferase assay reliability. Two versions of PTEN 3’UTRs were used, pmirGLO-3’UTR-PTEN long represent 3’UTR length obtained from UCSC genome browser and pmirGLO-3’UTR-PTEN short represents miRDB miRNA target prediction program version of 3’UTR length. A) pmirGLO-3’UTR-PTEN long and B) pmirGLO-3’UTR-PTEN short (*p < 0.05; Student t-test).

**Supplementary Table IA:** RT-qPCR primer table

**Supplementary Table IB:** Primers for amplifying 3’UTR sequences and restriction enzymes used for pmirGLO-Dual luciferase vector cloning. With red is marked restriction enzyme cutting sequences and with lowercase letters 5’overhang sequence

**Supplementary Table IIA**: Affymetrix microarray results upon KGN cell transfection with hsa-miR-548ba, adjusted p-value <0.01

**Supplementary Table IIB**: Potential target genes of hsa-miR-548ba. Overlapping part of Affymetrix microarray results (asjusted p-value <0.01) and bioinformatically predicted target genes (predicted by at least 2 programs out of 4). Bioinformatical programs: DIANA microT v3.0, microT CDS v5.0, TargetScan 7.1 and miRDB. Genes marked red were validated by RT-qPCR.

**Supplementary table IIC:** Affymetrix microarray results upon KGN cell transfection with hsa-miR-7973, adjusted p-value <0.01.

**Supplementary Table IID:** Potential target genes of hsa-miR-7973. Overlapping part of Affymetrix microarray results (adjusted p-value <0.01) and bioinformatically predicted target genes (predicted by at least 2 programs out of 4). Bioinformatical programs: DIANA microT v3.0, microT CDS v5.0, TargetScan 7.1 and miRDB. Genes marked red were validated by RT-qPCR.

**Supplementary Table IIIA:** Reactome pathways regulated upon transfection of KGN cell-line with hsa-miR-548ba mimic. Pathways in bold are unique to hsa-miR-548ba regulation in comparison to samples expressing hsa-miR-7973.

**Supplementary table IIIB:** Reactome pathways regulated upon transfection of KGN cell-line with hsa-miR-7973 mimic. Pathways in bold are unique to hsa-miR-7973 regulation in comparison to samples expressing hsa-miR-548ba.

**Supplementary Material.** Target gene 3’UTR sequences and bioinformatically predicted target sites for miRNAs hsa-miR-548ba or hsa-miR-7973. Four target prediction programs were used: DIANA microT v 3.0, microT CDS v5.0, TargetScan 7.1 and miRDB. Red underlined sequences represent seed sequences predicted by both TargetScan and miRDB, green underlined sequences represent microT CDS v.5.0 predicted target sequences, blue underlined sequences represent only TargetScan predicted seed sequence location and yellow marks seed sequences predicted only by miRDB. Hsa-miR-548ba potential target gene 3’UTRs are: a) BCL2L11, b) LIFR, c) NEO1, d) PTEN, e) RARB and f) SP110. Hsa-miR-7973 potential target gene 3’UTRs are: g) ADAM19, h) ATHL1 i) ATP6V1A, j) FMNL3 and k) PXDN. Potential target gene for both hsa-miR-548ba and hsa-miR-7973 3’UTR is: k) TGFBR2.

## Notes

† **Grant support**: This study was supported by Estonian Ministry of Education and Research (grant IUT34-16); Enterprise Estonia (grant EU48695); the EU-FP7 Eurostars program (grant NOTED, EU41564); the EU-FP7 Marie Curie Industry-Academia Partnerships and Pathways (IAPP, grant SARM, EU324509); Horizon 2020 innovation program (WIDENLIFE, EU692065) and MSCA-RISE-2015 project MOMENDO (grant no 691058). Affymetrix microarray data is available at Gene Expression Omnibus repository, accession number GSE122731

