## Supplementary material for "MicroRNAs expressed from FSHR and aromatase genes target important ovarian functions": Supplemetary figures 1,2,3

### SUPPLEMENTARY RESULTS

**Supplementary Figure 1.** Transient hsa-miR-548ba and hsa-miR-7973 expression in KGN cell-line compared to primary human granulosa cells. For positive control human mural (MGC) and cumulus granulosa cell (CGC) RNA was used (n=8). miRNA expression in KGN cells is shown as average expression of four parallel samples ±SD on $log_2$ scale. miRNA expression levels were normalized against miRNA hsa-miR-132-3p levels. Transfected KGN samples are indicated with + sign in the legend (yellow and pink colour).

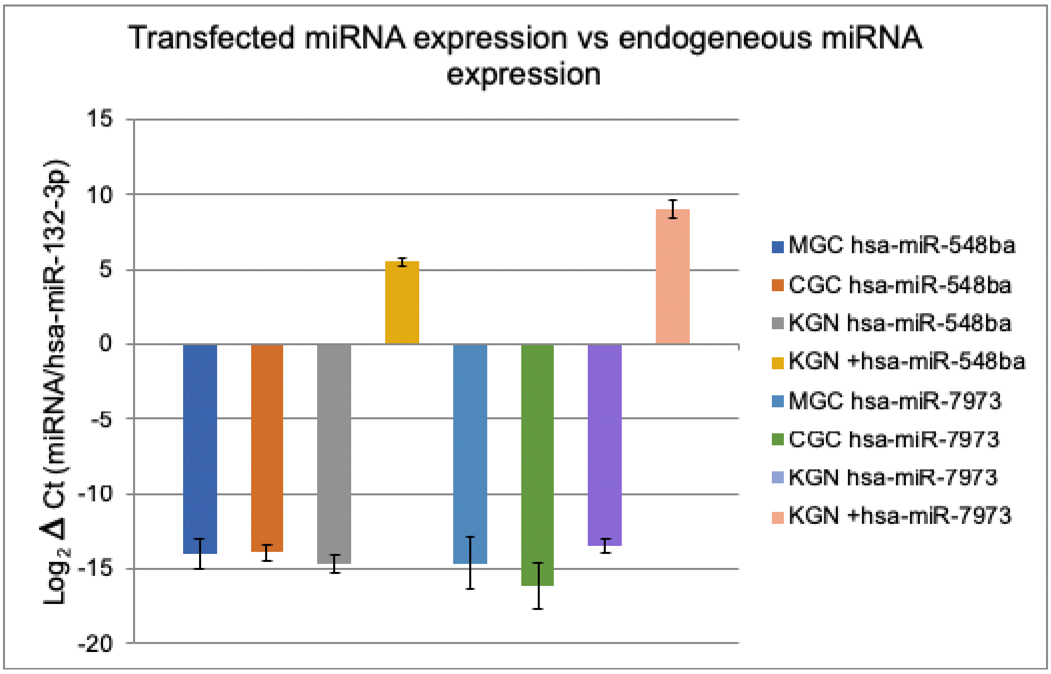

**Supplementary Figure 2.** KGN cell viability upon transient miRNA expression. Luminescence signal was measured from cells transfected with hsa-miR-548ba, hsa-miR-7973 or control miRNA cel-miR-39-3p at three time-points: 24, 48 and 72 hours. Signal from non-transfected cells (control) were measured at 0, 24, 48 and 72 hours. The indicated value refers to normalized luminescence: signal from all lysed cells minus signal from dead cells. Results are shown as average of 3 independent experiment with ± SD. Student t-test p-values were above 0.05, no significance difference in living cell number were detected between non-transfected control samples and transfected cells.

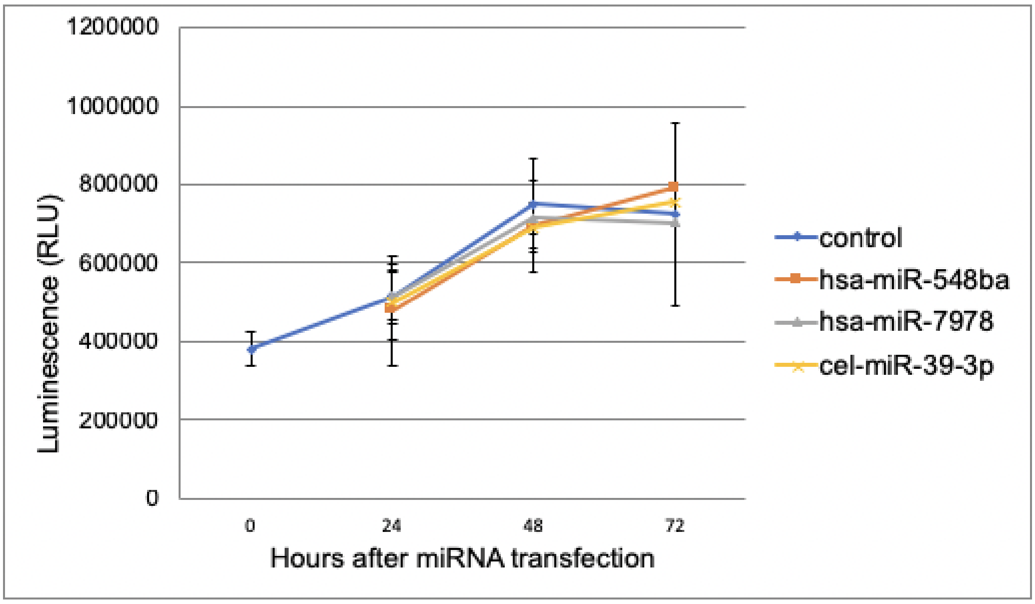

**Supplementary Figure 3.** Suppression of luciferase signal upon hsa-miR-21-5p binding to PTEN 3'UTR. pmirGLO-3'UTR-PTEN vector and hsa-miR-21-5p were used as a positive control for miRNA:mRNA binding to confirm the luciferase assay reliability. Two versions of PTEN 3'UTRs were used, pmirGLO-3'UTR-PTEN long represent 3'UTR length obtained from UCSC genome browser and pmirGLO-3'UTR-PTEN short represents miRDB miRNA target prediction program version of 3'UTR length. A) pmirGLO-3'UTR-PTEN long and B) pmirGLO-3'UTR-PTEN short (*$p < 0.05$; Student's t-test).

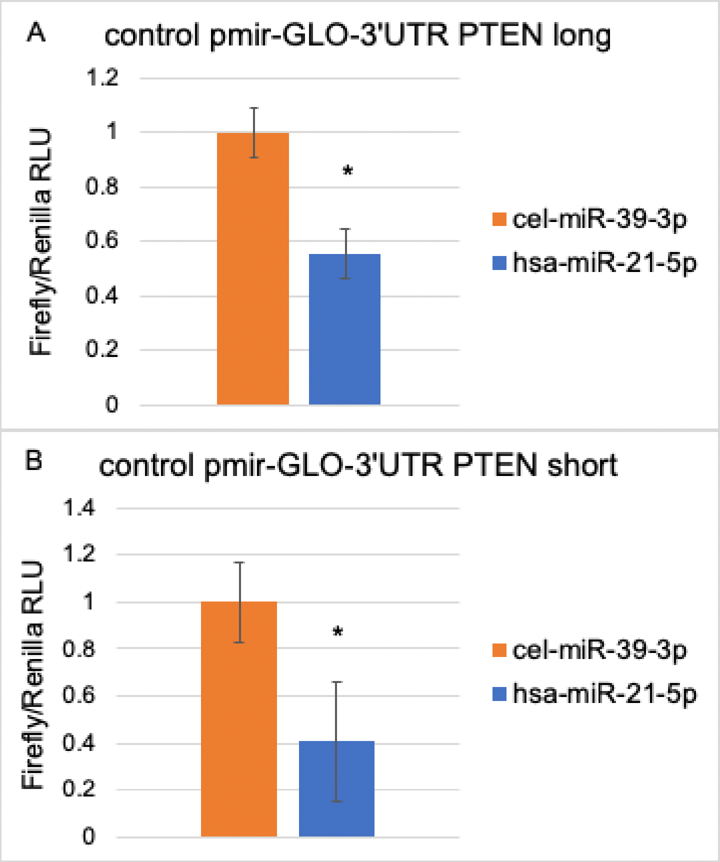
