## Supplementary Material for "MicroRNAs expressed from FSHR and aromatase genes target important ovarian functions"

**Supplementary Material.** Target gene 3'UTR sequences and bioinformatically predicted target sites for miRNAs hsa-miR-548ba or hsa-miR-7973. Four target prediction programs were used: DIANA microT v 3.0, microT CDS v5.0, TargetScan 7.1 and miRDB. Red underlined sequences represent seed sequences predicted by both TargetScan and miRDB, green underlined sequences represent microT CDS v.5.0 predicted target sequences, blue underlined sequences represent only TargetScan predicted seed sequence location and yellow marks seed sequences predicted only by miRDB. Hsa-miR-548ba potential target gene 3'UTRs are: a) *BCL2L11*, b) *LIFR*, c) *NEO1*, d) *PTEN*, e) *RARB* and f) *SP110*. Hsa-miR-7973 potential target gene 3'UTRs are: g) *ADAM19*, h) *ATHL1* i) *ATP6V1A*, j) *FMNL3* and k) *PXDN*. Potential target gene for both hsa-miR-548ba and hsa-miR-7973 3'UTR is: k) *TGFBR2*.

**a.** *BCL2L11* 3'UTR sequence with underlined bioinformatically predicted hsa-miR-548ba target sites. microT CDS v5.0 predicted target sites are marked in green: positions 144-163, 656-675, 2310-2328 and 2696-2717. TargetScan and miRDB commonly predicted seed sequences are marked in red: positions 2322-2328 and 2712-2718.

1 CAGGTTCTTT GCGGAGCCGA GATACCATGC AGACATTTTG CTTGTTCAAA CCAACAAGAC
61 CCAGCACCGC GGTCTCCTGG TGCCATTATT ATGCAGCCAG CGGTTCTCTT GTGGAGGGGG
121 CAGGTGACGT TTCAGAAGAC ACCG**AGCTGG ATGGGACTAC CTT**TCTGTTC ATCACCACAC
181 AGCAGAATTT CTAATGGAAG TTTGTTGTGA ATGTAAAGGA GGGAGCATTC TTTGCTTTTT
241 AATATACAAA CCATGGTTTT TTGGAGCAGG ATTTTGTGTA AGAATGGTGT TTACATGCAG
301 TGTGTTTTCC CCCTCACCTT CAATAAGGTT TTTCAAAAAG GAAATGGAAA CTTTTTAACC
361 AATTTGTGAA TAACTTTTGT ATTAAAATTT TAAGAACCTA CGGCCTATTC TCAGAGGATT
421 ATGTAACCCC TGCAGTGGAA ACTGAGCCAG CTAACTTAAA AAGCTGCCTT AGTTTATTTT
481 TAGAGATTAC AGAATTTTTA AACAGGGAGA CGTGTGATAT ACTCCCTCCC TTCCCTACTA
541 TTGCCTCTCT GACCTTTTTA AATTATTTTT AATACCAAAA GAGTTCTTTT GAAATGGAAC
601 TGATTAAAAG GGCAGAGGGT CTGTTGCCAG CCTGCATTGA TATACCAGTC CCATTT**GTAA**
661 **ATATTTACGT ACCTT**TATAA ATTCAGTTGC ATCTGTGGCA AAATTTCAGA CTATTTTTGC
721 GTCTTTCCTC ATCACTTTTT GTGATGCAAC TCCAGTCTGG ACTCAGATGC ATAGATTTGG
781 TCCAGTGTAT TTTCATGATA AAGTGAAATT GAGTCAGAAC AAGAGTTAAT ATCTGCCTGT
841 ATCTTGCACA GTTCGAGCGA TCTGTTATTA ACTGGGAAGC ATTTGGTGTT GGTTTTCATT
901 CCATTTCGAC GAGCATGTTA TTGGGAAGTA TTCTGAAGAG GCAATAGCAG TAATAACAAC
961 AGACTTAAGT GCTACGCCCC TTTGTGCTGC TGGCTTTTCT GGTTGCAGGC TTTCCCATGG
1021 TCACAGGATG CACTGTCAGC ATCAGGTCCC AGAGGGCCAC CGTGTCCATT ACAGCAGAGT
1081 CCAGCTGCAG CATCCAGCTC ACGCCCTCAT GGGAATTGGC ACAGGCCTGG GGCAGGGCTT
1141 CTGATGGCCA TTTGCTTGGC CTCCTGCATT TTAGTCCAAC TCACAGTCCA CTAGCTTCAC
1201 TCCTTTAAAT TCACTTTGAA ACAGGCCTCA TCCCACTTCC ACCAGCACCA TAGAAGAATA
1261 ATTCTGGGCA GAAGTCTGTT TTTTTTCATT TTTCCAGGAC AGTTGGATAT TGTCAGGCCA
1321 CTTGTGACCC CAGCCATGTA GTGAGGGTGC TCTTTCTCTG TGCCTGCTCC TTATGAGTGC
1381 AGTGGAAGGA AGCCACACAC TGGTCAGTCA TTTCAGAGGC AGCAGATGCC CAGGGAGACC
1441 CAAGAAAGAG TCAGGTTAGG GAGCAGTGAA AGTGAGGAGG GAAGACAATT CTGTGAACTC
1501 TGTAACTCTT AAAATTTTTG AAAACTCCAT CGTTAAACAA CTTTTAAAAG AAATAACTAA
1561 ATTTTCAAAT GAGTAAGCAG TGCCACCAAC TAGTGTTTTG CCCGATAGAA GAGCCAGCAT
1621 GTTCACGTTA TTTAAATTAG GTGGAAAAAT CTAAACATTT TTATCTTCAT AATTTAAAAA
1681 ATATATATGT ATATATTGCA TATTCACTTT TTCCTTTAGG TAGAGATGAT TTCAATCCAA
1741 ATACTCTTAC TTTAAAAAAT TTCCTTTCCC CAAGAATCTC CTTGGGACTT TGACTTATTT
1801 TTAAAGCTGT GTTGGAGCTC ATCTTGTTCC CTGATGTGTC TCGAGCCCAT TGGTAGGGTC
1861 ATACAAAGCC CACGGTTACA AGCAGTGGTA GGATTGCAGC CGTGGGCCTG CTGGACACAC
1921 ACATACACCA AAGATGTATT TGGATCTGGG CACCCCCTCC CAGGATCCCT GTACTCACGT
1981 GCCAGTCTCC TGACTAGAGC ACTTTACTCT GTTTCCTCAG CCCTGCAGCC CCTGGGAGCA
2041 CACACTGGGT GCAGCCCTGG GCCAGGCACG GGAGGCCCTG CCCTGTGCTG CCCAGGGGCT
2101 GTGTGCACCA CATGAGCACA TTTCCCTCTG GCCTGGCGGC CTCCAGGCTG GCTGTGGAAA
2161 CAGTTCCTGA GGAAATTAGA GATTCTATGA ATTGTAGGAG TATTAAAGAC CAGGCTGTTG
2221 GCACCAGAAC TTAAAGCGAT GACTGGATGT CTCTGTACTG TATGTATCTG GTTATCAAGA
2281 TGCCTCTGTG CAGAAAGTAT GCCTCCCGTG **GGTATACGTT TTTACCTT**TT TTAAAAAACA
2341 TTTTTGTAGA AAAAATAATT AAATCCCCTT TTTGGAAACT TACTGCAGGT TTTGTGCCTT

2401 GACAACCTCT CCCTATGTGA GGTTTGTAAA AAGTGTCCTG TGACTTAACA CAGAAACGCA
2461 ATAAACACAC ACAAAATAGT TTCATGAGTG ATTCTTCAGA TGCCCTTCCC AACTGGTTAG
2521 TTGATCAAGA ATTTTGGGGG TGGGGGTTGC GGAGAAATCA AGTTTAAAAT TCCTTCTGAT
2581 TAAAAAAATA TAGTGGAATA CAATTGTCTG CCGTTTCCCC TTCTTAATGT ATATATTGTG
2641 AGTATTTATT AGATTCGTAG GTCATATTAC TTATCAACTG AGCCAAATGT CTGTGT**GCAA**
2701 **TTGTGTTTCC TTTACCTT**GT AAAATTTTGT ACAGCATAAA TAAGTAAAAA AATCACTGTT
2761 TTTCTCAACT TTTTCAAAAT CAAGGATTGT AAATATTGTA GATTCTTTTT CTGTGTGATG
2821 TGTCCTACTG TTTCATAATG CTGTAACTTG TAGAAATATT GTATATTTAT TTTCTGCTTA
2881 TTTAATGTCT TAATTTCTGA AAAGTATTAA CATCCCTGTC TCCCACTCCC CTGCCGTCCC
2941 ATGAAGTTAA CTCCTGAGAG TTGTCGGGGG TGACTGGAGA GCTCATTGCA GACCACGTGG
3001 TCCTCCAGGG TGGCTCTCCA CCTTCGGGTC CTGGTATTTC CAGTCAAGTG GGTTTCAATT
3061 CTTGGGCTTT GCCGCCCTTA TGATGAAGTG TGTGTTTGAT GCCAGTGAGA AACTCAGTCT
3121 GGCAGGCTAC AAAATTCTAC TCCAAGAAAT ACCCAGCAAC CTTCTGTTTG TTCCAAAGCA
3181 ACTAGCTTAT CATGCAAGCA AATTTTGCTG ACTCCAGGCT TTATCTTTAG GAAAACAAAA
3241 AAACCAAAGT ATTATCAGCA GGTGGGAAAG ATTTTTCTAT TGAAAATTTA TCCCTGACAA
3301 CTCAGCGTTT AGAAAAGAAA TAAAATGTGC CACTTCCAGA GGTGCTGCAT TGCAGTTGTT
3361 CAGGGCTAGG GCCAGGCAGG ACAAGTGAAT GGGTGGGACA GGTGGCTCCT GCCTAAGGAC
3421 CACCTCAGGC CACTAACCCC TTGTGGACAA CTGTGAGTAG CTGGGTTTTC CCCCACCTGC
3481 TGTGCAACTT CCTGTGCTTT GAGGTTGGAC TAACTTGTCT TCAGGAGCTA ATTAACTGTA
3541 CAGCCCTCCC CACGCCCCAC CCATACGGTC ACTGCATTTG GTCAGCCTGC TTCTTCAGGT
3601 CGATGCCCTC CTTCTGATAC TCCATCTCCT TCAGGGGAGG TTGGGGCCCC ACTGGACTGG
3661 GTGTCAAGAT GTGAAAGCTT ATGGGAGCTT TAAGGAGACT TCATGGTGGT TCCATGCAGG
3721 TGGTTCTGCC ATCCCTGCTG ATTTAGCCTG GTGCCTGTGT GTGTCCACTC ACGTACACGT
3781 GGGGTGGGGG AAACGTGTCT ACAGATGACG CTAAATCAGT TGGGGTCTAC TCTAAACAGC
3841 ATTGTGTGTA AGAAGCATCC TCAAGCTCCC AGTTAAGTAA CTTGACTACT TTTATTTGGG
3901 AATTTCAGAC TATAGAAGCT CTCTTATGTT TTATGTCCAG ATTCTGTGAC CACTAGTTAC
3961 TGTATCAGAA CTCATCAGGT ACCCACTTAT AAATAGCACT GATCTGGCTG TATACTGATC
4021 CATCACTAAC CTGTTTTCTA GGACCCAGCG TATGTAGCAT TTGTATTGCA GTTTCCCTGG
4081 CTTACTTGTG TTTTGCACTG ATGAATTTTG ACAGGGTAAT TGCCACTTTA CTTGTGCAAT
4141 ACTGCTGTAA ATAACTGCAG ATTTTTAAAC AATCTTTTAT GTTAATTTTA TAAAAATAAA
4201 ACTTTCAACT AGTTAAAAAA A

**b.** *LIFR* 3'UTR sequence with underlined bioinformatically predicted hsa-miR-548ba target sites. TargetScan predicted seed sequence locations are marked in blue: positions 1384-1390 and 5865-5871. With green are market microT CDS v5.0 predicted target sequences locations are marked in green: positions 760-777, 1064-1092, 4378-4397, 5241-5267 and 6366-6391.

1 CAGTGTCACC GTGTCACTTC AGTCAGCCAT CTCAATAAGC TCTTACTGCT AGTGTTGCTA
61 CATCAGCACT GGGCATTCTT GGAGGGATCC TGTGAAGTAT TGTTAGGAGG TGAACTTCAC
121 TACATGTTAA GTTACACTGA AAGTGTTCAT GTGCTTTTAA TGTAGTCTAA AAGCCAAAGT
181 ATAGTGACTC AGAATCCTCA ATCCACAAAA CTCAAGATTG GGAGCTCTTT GTGATCAAGC
241 CAAAGAATTC TCATGTACTC TACCTTCAAG AAGCATTTCA AGGCTAATAC CTACTTGTAC
301 GTACATGTAA AACAAATCCC GCCGCAACTG TTTTCTGTTC TGTTGTTTGT GGTTTTCTCA
361 TGTGTATACT TGGTGGAATT GTAAGTGGAT TTGCAGGCCA GGGAGAAAAT GTCCAAGTAA
421 CAGGTGAAGT TTATTTGCCT GACGTTTACT CCTTTCTAGA TGAAAACCAA GCACAGATTT
481 TAAAACTTCT AAGATTATTC TCCTCTATCC ACAGCATTCA CAAAAATTAA TATAATTTTT
541 AATGTAGTGA CAGCGATTTA GTGTTTTGTT TGATAAAGTA TGCTTATTTC TGTGCCTACT
601 GTATAATGGT TATCAAACAG TTGTCTCAGG GGTACAAACT TTGAAAACAA GTGTGACACT
661 GACCAGCCCA AATCATAATC ATGTTTTCTT GCTGTGATAG GTTTTGCAAG CCTTTTCATT
721 ATTTTTTAGC TTTTATGCTT GCTTCCATTA TTTCAGTTG**G TTGCCCTAAT ATTTAAA**ATT
781 TACACTTCTA AGACTAGAGA CCCACATTTT TTAAAAATCA TTTTATTTTG TGATACAGTG
841 ACAGCTTTAT ATGAGCAAAT TCAATATTAT TCATAAGCAT GTAATTCCAG TGACTTACTA
901 TGTGAGATGA CTACTAAGCA ATATCTAGCA GCGTTAGCTG TCCATATAGT TCTGATTGGA
961 TTTCGTTCCT CCTGAGGAGA CCATGCCGTT GAGCTTGGCT ACCCAGGCAG TGGTGATCTT
1021 TGACACCTTC TGGTGGATGT TCCTCCCACT CATGAGTCTT TTC**ATCATGC CACATTATCT**
1081 **GATCCAGTCC TC**ACATTTTT AAATATAAAA CTAAAGAGAG AATGCTTCTT ACAGGAACAG
1141 TTACCCAAGG GCTGTTTCTT AGTAACTGTC ATAAACTGAT CTGAATCCAT GGGCATACCT
1201 GTGTTGCAGG TGCAGCAATT GCTTGGTGAG CTGTGCAGAA TTGATTGCCT TCAGCACAGC
1261 ATCCTCTGCC CACCCTTGTT TCTCATAAGC GATGTCTGGA GTGATTGTGG TTCTTGGAAA
1321 AGCAGAAGGA AAAACTAAAA AGTGTATCTT GTATTTTCCC TGCCCTCAGG TTGCCTATGT

1381 ATT**TTACCTT** TTCATATTTA AGGCAAAAGT ACTTGAAAAT TTTAAGTGTC CGAATAAGAT
1441 ATGTCTTTTT TGTTTGTTTT TTTTGGTTGT TTGTTTGTTT TTTATCATCT GAGATTCTGT
1501 AATGTATTTG CAAATAATGG ATCAATTAAT TTTTTTTGAA GCTCATATTG TATCTTTTTA
1561 AAAACCATGT TGTGGAAAAA AGCCAGAGTG ACAAGTGACA AAATCTATTT AGGTTCTCTG
1621 TGTATGAATC CTGATTTTAA CTGCTAGGAT TCAGCTAAAT TTCTGAGCTT TATGATCTGT
1681 GGAAATTTGG AATGAAATGC AATTCATTTT GTACATACAT AGTATATTAA AACTATATAA
1741 TAGTTCATAG AAATGTTCAG TAATGAAAAA TATATCCAAT CAGAGCCATC CCTAAAGAGT
1801 GTTCTCTTGT CTTCCTTTGT ATCCTCTTTG GCTCCTTCCC TTAGCTTCCA CCCCAGACAT
1861 CAGACCTGGG GCTGCCTTCT CTCCCTGCCA AAGCTTGCTG TTAGCATTGT CTGCTTGCCC
1921 TCAACTCCTG CCTTGTTCCC TGTTGAAATG TATCAATCCC GTACTTTATC TCTCCAGGAC
1981 AAGGATAAGT GTGGCTACAG TAAAAGTTTA TTGCTCTAAA CGGCGCTAAC CTGAGATAAT
2041 GAGAAAAGGC CAGAAGAGAT TGTTTGATCT CTGTCTGTCA ATATATATGC ATTTCAACTG
2101 TGACCATCTG GATCCAGGGC CCTTTCTAAC CTATATAACG ACTCATTTGT TTGGTTAATG
2161 ATGAATATCA CTGGGGCTCT TAAGTCCTTT CGTTCTATCC AAGACTTCAT TGTTTTTTTC
2221 TGTATGAAAA AAGTGCAGAA TATCACTCAC CTTTCCCCTA CTTTTGGATC CCCTACACGA
2281 GGATCACTAG CTTAATGAGA AGCAGAGTAA CGCTATTAAG TCTGGGCAAG CTCCTTATAA
2341 GTTGCTGTGT GCTTAAAAGA ATGTGATTTA TGTCTGGGGC CAAAGCAGTT TTCCTTCTTT
2401 TCCCTCATTG AATTCAAGAC AGCTAGACCC TTGTGTAGAT CCCAGGATAT TACCTCTACT
2461 CCCCCCAACC TGTGTGTGTA GGGAGAGTAA CAACACGGGA AGAGGGGAGG GAAGTTGGAA
2521 GGAGATTGGT CACATGTTCA GTTGTCTGAG ATACTGGTAA ATGTGCGCTG TTTGTCAATG
2581 AAGGTTGCAT GCCTGCATTT ACAAGGTGGT CTCATACTGA TTGTGCTCTG TCCAGTTAGA
2641 TTCGTGGCTT CTCCCTTGGA AGCTACAATT AAGGACTAGT TAGAAGCAGA GGCTAGATGT
2701 GTTCTCTCCA GCCCTTACTA ACTGGGAAAA TAATTGAGAA ATCCCTCTTT TCCTCAGTGA
2761 AAGCCCAGTC ATTGGTTAGA AAAATCTAAT TGTCAAACTA GCATTATGTT CCATCATATC
2821 TGTGGGATAG TAATTCATGT AAGACACATT TTTACCTCTC TTACAAGTCA ATTCACTTAC
2881 AAGCCAGTTT CAAATCAAGC TTCCCTTGCG AGGAATGGAG ATCATTATGG GGCTTATTAT
2941 GACTACATTT CTTTTATGAG TGACCATTTT AAATAATTAA TATTTTATTT AATATTAAAT
3001 TATTAAAGAC TTAGCTAGAC TTTTAACATT AGATGAACCT TGTGAATCAT ATATTGCATA
3061 GATTAGCACA TGTATGCAAT CTAGGTCATC AATTTGTTGA ATTTATACAG CCCTAGATTT
3121 GAGATTGTGG CCCTGAGTAT CTGGATAGTT GAGTTGTGTG TTTATGTGCT TATATGATAG
3181 AGGTGCTTTG TTCCTGCAGC TTAGAAACAG AAGCAGCCAT TCACTGAGAA GCTTAAGAGA
3241 GGAAGTATCA ACCTCATTTT CCAAACCAGC ATTACTTTCA CCGCTCGTGG ATCTGAAAGA
3301 TTACATTTAG AACATTTAGT ACTATTACAC TCTCAGAAAC TGTGGTAAAA AGTCACATTT
3361 TCAAAACTTC ATGCATATTG TATTCTTGTT GGAAATAAGT CCTATAGTTT CTTAATTGTC
3421 TTCATGCCGT CCATATTTAA CAAACATTGC AAGTCCTTTT TATATTTGGG TAATTATTTG
3481 TAATTTAGTG AAGGAGACTA GGGGATGTTT TCTTCCAAAG GGAATTTAAA ATCAATTTTA
3541 TGGTATTTTG AAAGTAAAAT ACTCCTTAAC GTGTCAATAT TTTTAAAATG TTATCTCGAA
3601 GTTTTCTATT GCTCAGGTAT ATAATCTAAT AAAATTTTTT GTTTGCTCTA AGGTTATTTT
3661 TGCTATTCTC TATTTTGAAT TATGTTAAAA TTTGTGTGCA TTTTATGAAA TGCCTTTTTT
3721 CTTATTTTGA AAGTGTAAAA TTGGGCATCT CAGAGTAAAA TGTGGGATTA TTAGCTACAA
3781 AGACTAGGCA CATCGTGAAA GCTCAGTAAC TAATTTTCTT GACTGAGTCC TATCCCAGAG
3841 TGTCACAGAT TCAACAGAAT TTTCCTGAAG GGTGATACAC ACCAAAATCT AGCTTGTGTG
3901 GATCTGAGAC TATTGACTCA AAGATTCAAA TCTACTTCGG AAGTGACTCC ACGGTTTGAG
3961 TGAAAACTGA GGAAGGACTC ATGCCTGTAT ATCACTAGAT ATCTTATTTC TCCATAATTT
4021 TAAGTGTTCA TAATTTCTTC TTCTTTGATG GGTGTGCAAC TGGGGCATGT TTGTACCAGC
4081 CAGCATGTGT TCTGCCCAAC AGTGTGTCCG TGTGTGAGGT TAACTGATTC AGAACTCATT
4141 CTCTTGATAA ACTATTTCTT GAAAGCATTT ATTTTAAGTC TTTATTTCCT GCTGAAATGA
4201 AGTTTCATGC TGTCATGCCC ATCTTATTTA TTCTAGAAGA AAATTTTCAT AGAAGAAAAT
4261 GATTTATGGC CAAAGTTACA GAACATTCAT CTTTTAGACT TAGCCTTAGA ATATCATTGG
4321 AAATTACTTG GCTTGAATTT GGTTCGTTTT TATCTTTATG GATTATGTAC CAGGGCT**ATT**
4381 **AATAATACTT AAACCCC**TTA TTTTGCTATG GTGGTTAGAT TTTTTTATTC CTTTTCAATT
4441 AGTGCTATTA ATTATTACAT GGTCGTATTC ATGCTTATTA CAGATGAACT ATTTTTAGTG
4501 CCAGTGTAAA GGATTAAGAA TAACATTCAA TTAGTTGGCT ATTTCAGGCT TTATTGTACA
4561 TGGTAATGTT TCAGGGAGTG TGATTTGGTC TAGAATATAT CCAGATCCTT GTAATTATGA
4621 TATTGGGTTA TGACAGGATT CTATTATTTT TAGCTATGAA TGGTGCTTAT TTTTCCCTAT
4681 ATTAGGAATG TTAATATTTG GATTATCAGA GTCCATGTTA TAAACAGAAT TTACCTATGC
4741 AAGACAAATT CTAACTGTGC AACATTTTCA TGAGAATTAA AAGTTGTGTG TTAAGGGGAA
4801 AACCCTCTGA ATGTTTTTAT GTGTAAGTGC CTAAAGAACC GTAGAAACAG TACATCAAAA
4861 GCTCTTATAA AAGAAAAACT GATTGAACAT TAGCCAGAAA CAAAATATGA TTTACATTTT
4921 TTTTTTTGGT TGCTGGGACT GAATACCCTA AAAACTAAGA GATGTGAGTG GGGAAAATGT
4981 CTAATTGATA AGAAAGTTGA TTTATATTAT ATTTACATGC ATCTTCTAAG ATTGCCTTTA

5041 CAGTAGCCAA TCAGAACTTT AAAAAAAAAA AAGCTAACAA CATAGATCCT ATGTGTTGCC
5101 CTTAAGACTA AAATACTTTG CACACGTGAT TGAATGGCTC ACTTAAGTTG TTGAATTTCC
5161 TTATACATTT AAAGTGTACT TCCGTTAAGA ATTATTCCAT TTTAAATGGG TAACTTTCAG
5221 CTTCCGAACT ATTTTCCTAA **ACATTCCAGT TGTGTATTAT AAAAGAT**CAA AATATCTCTA
5281 GGTGGTAGTA CAATTTTCAT TTATAACGTG GAAAAGTTAA CTGTTAGGTT TAACATTAGT
5341 ATTAGTGTTT TTGCAATTGC TCATAGAAAA GAACACTGAA TAATGGTTTT CATACAATTT
5401 TTGTAAAACA TTCTTCTCCC AGTTTGATTC CAAATGCTTT TCCCCAAATT GTCTAATATA
5461 TCAT**AAGCTA G**TGTTTCCTT GTATTTAATT TTTCCTTCCA ACACAATATT TAGGAAAAAT
5521 ATGAATAACA TTGTGAGTAG CGGGAGACAA CACGAAAAAG AAACCTTTCA TTGTTGTGGA
5581 GGGGTAGGTG TTATGTGATT GATGCCCTTC CTGGCGTTTG TTTTATCCTG TTATTTTCTC
5641 AAAAAAGAGC ACGGCTTTTT CTTAAATACA TTTAAATTCA GAATTGCTTG CAAAAGATTG
5701 GCCTGGATAG ATAGACATCC TAGATTTAAG ATAGTACCAT CTTAAAAGTA TGTTGAAAGG
5761 TAAAATAAAT CATTTCAAAA TACTTAACTT TCCTAATAAG GATGTTAGGT TTTCTTCTCT
5821 TTCATTAGCT GGCCATAAGT CAAGGTCCTT ATAACACTAG GAAATTACCT TTTTTTACTT
5881 TCAAAATTGC ATTATAGTGA ATTACTGTTT ACAAAATACA CATACTGTGA ACAGATATAA
5941 GTTTCTGTCT TTTATAAGGT TTGTAGAATG AATGTTTTTT TAAAGTGAGT ATTATATGTT
6001 AAATATTTTA AGTTTTTTAT AAATACTAGT AACTGTTTAC TAATTTTTGT TTGGTCAAAT
6061 GCTTGTAAAT GTAGCTGAAA GAAGATAGGG AGAAACTGCG GATCCCAAAC TGTTCCCTTT
6121 TTCATTTCTT GAAATGTTAC CACTACAGAC ATTTTTTTAA GGTGAATAAA CAGTTGTGAT
6181 GTGCTGTACC TAAAATCATG TTTAATCGTA TAAGGAAACA TTTCAATACA CTTATACAGG
6241 AAGAAAACTA TAGATGAAGT ACATGTGTGT GATTCAGTCT GATTCACAGA ATTCTGAGAG
6301 TAATATGGAA TAAAACAACT CCACTTAGAT GATAACTGAA GCATTTCCTG CCTTGTGAAA
6361 AT**TTGGATTT TAAATTGCTG TTAGAATGGG A**AATTTGGAC ACTTTATATC ATTGTATAAT
6421 TTCAGAATTT AGTTTCTGTA TCTTTTGGAA AACATGATTA TAGCAAAAAC ATAGAAAATA
6481 ATCTATTACT AAAACACCAT AAATGTAAAA CTAGTATGCT TGGCTGTTAA CTCTAAAGAT
6541 GTTACTTATG TCTGTTTTTA AAACATGCAT GTATTTAACA ATTTTATCAT AGTATTGTCA
6601 TGGAAAAAAT AAATATATTT TCTTACATCT TA

**c.** *NEO1* 3'UTR sequence with underlined bioinformatically predicted hsa-miR-548ba target sites. microT CDS v5.0 predicted target sequences locations are marked in green: positions 327-346, 390-407, 1982-2007 and 2190-2211. TargetScan and miRDB commonly predicted seed sequence location is marked in red: position 402-408.

1 CGACCTTCAC CAGGACCTGA CTTCAAACCT GAGTCTGGAA GTCTTGGAAC TTACCCTTGA
61 AAACAAGGAA TTGTACAGAG TACGAGAGGA CAGCACTTGA GAACACAGAA TGAGCCAGCA
121 GACTGGCCAG CGCCTCTGTG TAGGGCTGGC TCCAGGCATG GCCACCTGCC TTCCCCTGGT
181 CAGCCTGGAA GAAGCCTGTG TCGAGGCAGC TTCCCTTTGC CTGCTGATAT TCTGCAGGAC
241 TGGGCACCAT GGGCCAAAAT TTTGTGTCCA GGGAAGAGGC GAGAAGTGCA ACCTGCATTT
301 CACTTTGTGG TCAGGCCGTG TCTTTG**TGCT GTGACTGCAT CACCTT**TATG GAGTGTAGAC
361 ATTGGCATTT ATGTACAATT TTATTTGTG**T CTTATTTTAT TTTACCTT**CA AAAACAAAAA
421 CGCCATCCAA AACCAAGGAA GTCCTTGGTG TTCTCCACAA GTGGTTGACA TTTGACTGCT
481 TGTTCCAATT ATGTATGGAA AGTCTTTGAC AGTGTGGGTC GTTCCTGGGG TTGGCTTGTT
541 TTTTGGTTTC ATTTTTATTT TTTAATTCTG AGTCATTGCA TCCTCTACCA GCTGTTAATC
601 CATCACTCTG AGGGGGAGGA AATGTTGCAT TGCTGTTTGT AAGCTTTTTT TATTATTTTT
661 TTATTATAAT TATTAAAGGC CTGACTCTTT CCTCTCATCA CTGTGAGATT ACAGATCTAT
721 TTGAATTGAA TGAAATGTAA CATTGAAAAG ACTTGTTTGT TGCTTTCTGT GCAGTTTCAG
781 TATTGGGGCG GGTGGGGGGC TGGGGGTTGG TAATAGGAAA TGGAGGGGCT GCTGAGGTCC
841 TGTGAATGTT TCTGTCATTG TACTTTCTTC CAGAAGCCTG CAGAGAATGG AAGCATCTTC
901 TTTATTGTCC TTTCCTGGCA TGTCCATCCT TATTGTCACT ACGTTGCAAC TGGAGTTTGA
961 TTTGGATCTG GTTTTAAAAT TCTTCTGTGC AATAGATGGG TTTGAGGATT TAGCGGCCCT
1021 GATGTCTTGG TCATAGCCTG GTAAGAATGT CCATGCTGAG GAGCCAGATG TTGTATTTCT
1081 AACTGCCTGA GTCACACAGA ATAGGGTAAG AGCCTGACCC CATTCTGTAA ATCAGAAAGC
1141 AAGGATGGAG ACCCTTTCCT GCTGCTATTA TTGGCTCTCT TTGAGGAAGT TGGAGGTTAA
1201 GGAAGGAACT TGTTTGTTTC CGTATACGAC TCCTTCTTCT CTCTAGTTCA GTCTTCAGCC
1261 AGTCCAGCGC TCTCTTCCAC ACTTCAGAGC CCCTTCAGAG AAAGCATTAG CAGGAATGAG
1321 ACAAGGCAGA GCTGCAGTGC CCCCTGAGGC TTCCACACAT CTTTCTGAAT ATTATTTTTC
1381 AAGTAACAAG GGCAGGGACA GCGGAAACAG CTGCCCACCC CCCCCATCCC AGCAGCTCAG
1441 CTAAGCCCTG ATGAGAATGA AGCCACAGGA GTTGTCTGAG GTGAACCCAG CCGCTCAGCC
1501 ACACATGGAA GCCATTGCCT TTGCACATAG TTCTTGGGTT CTTTTTCCTA AAAAGGTAAG

1561 GAGCTGAGGT GTGTGGTTTT TTAATATTAA GAATATATAA TGGAAAACAC ACGACTGACG
1621 CTCAGGCATC TTCCCCTACT CCCCAACAGA TCCCCAGAAG ACAGCGTGGA AGGCAGTGTA
1681 GACAGTAAAT CGGGCTTCAG TTCTATAGCC AAGAAGAGAT CAGCTGCTGA AACCACCAGT
1741 GGGTACCCCA GGCCACCTGC CTTTGAACTT GGGGATTTGC CATGTTTGAT CTTGTCACAT
1801 ACTTGCTTTT TTACAAGATG AACTCTTTGT ATTTATGATT TGGGGGGCAA TGAAAGGTGC
1861 AATGCAGGAA CTGCTGCTGC CGAGCTCGCT GGTCACATGG GGGTGCCAGG CGGGATTCTG
1921 GAAAACCAGT GCACTTAAAC TGATCCTGAA GAGAGCTGTC CCAGCACTCT GGCCACCAGG
1981 A**GGGCCAGAT TCCCCAGAAA CTACCTT**TTG CCCAAAGAAC ATGCTCAGTA TTTGGGGCAT
2041 TTCCTCCCAC AAACCCTGAC TGCTTCTGTT ACCTCAGGGC CTTGGTACCT GGATACTGCC
2101 ACAGAATTGG GGCGGGTGGG GGAGGGGCCT ATTTTTAAAT AAAATAACTG TTCAAAGTTG
2161 GGGGTTTTTT AAAAAAATTAA GAAAAAGGA**A AGCTATTCTG TATTGCACCT T**TTCACAATT
2221 TAATACATTT TCTTACATTT TCCTGTGATT TTCGAAACTA AACCATTGTG TGTCCTGTAG
2281 TGTCCTGGTT GAGCTGCCGC TCAGCAGCTT CCTCGGGGGG ATTTGGAACA CCTGTGTCTG
2341 TCGCCGCACT GCCTGTGGGA GGGGCCCAGA GGGCTGCTGG GACTGGCGTC TGTACACACT
2401 TGTTTGGCCT TTTCTGTAGT TGATGCTGTA AACTCTATGG CTTTTTAAAA ACGATTTCAT
2461 GTTTTTATTT AGTATTGGAA ATCCAATACA CTTTTTTAAT CCAATCAAAC

**d.** *PTEN* 3'UTR sequence with underlined bioinformatically predicted hsa-miR-548ba target sites. microT CDS v5.0 predicted target sites are marked in green: positions 51-56, 202-224, 566-585, 1890-1906, 2893-2913 and 2986-3006. TargetScan and miRDB common predicted seed sequence location is marked in red: position 2907-2913. For PTEN two isoforms of 3'UTR sequences were used: PTEN short 1-3,300 and PTEN long 1-6,458 nt.

1 ATTTTTTTTT ATCAAGAGGG ATAAAACACC ATGAAAATAA ACTTGAATAA **ACTGAAA**ATG
61 GACCTTTTTT TTTTTAATGG CAATAGGACA TTGTGTCAGA TTACCAGTTA TAGGAACAAT
121 TCTCTTTTCC TGACCAATCT TGTTTTACCC TATACATCCA CAGGGTTTTG ACACTTGTTG
181 TCCAGTTGAA AAAAGGTTGT G**TAGCTGTGT CATG**TATATA CCTTTTTGTG TCAAAAGGAC
241 ATTTAAAATT CAATTAGGAT TAATAAAGAT GGCACTTTCC CGTTTTATTC CAGTTTTATA
301 AAAAGTGGAG ACAGACTGAT GTGTATACGT AGGAATTTTT TCCTTTTGTG TTCTGTCACC
361 AACTGAAGTG GCTAAAGAGC TTTGTGATAT ACTGGTTCAC ATCCTACCCC TTTGCACTTG
421 TGGCAACAGA TAAGTTTGCA GTTGGCTAAG AGAGGTTTCC GAAGGGTTTT GCTACATTCT
481 AATGCATGTA TTCGGGTTAG GGGAATGGAG GGAATGCTCA GAAAGGAAAT AATTTTATGC
541 TGGACTCTGG ACCATATACC ATCTCC**AGCT ATTTACACAC ACCTT**TCTTT AGCATGCTAC
601 AGTTATTAAT CTGGACATTC GAGGAATTGG CCGCTGTCAC TGCTTGTTGT TTGCGCATTT
661 TTTTTTAAAG CATATTGGTG CTAGAAAAGG CAGCTAAAGG AAGTGAATCT GTATTGGGGT
721 ACAGGAATGA ACCTTCTGCA ACATCTTAAG ATCCACAAAT GAAGGGATAT AAAAATAATG
781 TCATAGGTAA GAAACACAGC AACAATGACT TAACCATATA AATGTGGAGG CTATCAACAA
841 AGAATGGGCT TGAAACATTA TAAAAATTGA CAATGATTTA TTAAATATGT TTTCTCAATT
901 GTAACGACTT CTCCATCTCC TGTGTAATCA AGGCCAGTGC TAAAATTCAG ATGCTGTTAG
961 TACCTACATC AGTCAACAAC TTACACTTAT TTTACTAGTT TTCAATCATA ATACCTGCTG
1021 TGGATGCTTC ATGTGCTGCC TGCAAGCTTC TTTTTTCTCA TTAAATATAA AATATTTTGT
1081 AATGCTGCAC AGAAATTTTC AATTTGAGAT TCTACAGTAA GCGTTTTTTT TCTTTGAAGA
1141 TTTATGATGC ACTTATTCAA TAGCTGTCAG CCGTTCCACC CTTTTGACCT TACACATTCT
1201 ATTACAATGA ATTTTGCAGT TTTGCACATT TTTTAAATGT CATTAACTGT TAGGGAATTT
1261 TACTTGAATA CTGAATACAT ATAATGTTTA TATTAAAAAG GACATTTGTG TTAAAAAGGA
1321 AATTAGAGTT GCAGTAAACT TTCAATGCTG CACACAAAAA AAAGACATTT GATTTTTCAG
1381 TAGAAATTGT CCTACATGTG CTTTATTGAT TTGCTATTGA AAGAATAGGG TTTTTTTTTT
1441 TTTTTTTTTT TTTTTTTTTA AATGTGCAGT GTTGAATCAT TTCTTCATAG TGCTCCCCCG
1501 AGTTGGGACT AGGGCTTCAA TTTCACTTCT TAAAAAAAAT CATCATATAT TTGATATGCC
1561 CAGACTGCAT ACGATTTTAA GCGGAGTACA ACTACTATTG TAAAGCTAAT GTGAAGATAT
1621 TATTAAAAAG GTTTTTTTTT CCAGAAATTT GGTGTCTTCA AATTATACCT TCACCTTGAC
1681 ATTTGAATAT CCAGCCATTT TGTTTCTTAA TGGTATAAAA TTCCATTTTC AATAACTTAT
1741 TGGTGCTGAA ATTGTTCACT AGCTGTGGTC TGACCTAGTT AATTTACAAA TACAGATTGA
1801 ATAGGACCTA CTAGAGCAGC ATTTATAGAG TTTGATGGCA AATAGATTAG GCAGAACTTC
1861 ATCTAAAATA TTCTTAGTAA ATAATGTTGA **CACGTT**TTCC ATACCTTGTC AGTTTCATTC
1921 AACAATTTTT AAATTTTTAA CAAAGCTCTT AGGATTTACA CATTTATATT TAAACATTGA
1981 TATATAGAGT ATTGATTGAT TGCTCATAAG TTAAATTGGT AAAGTTAGAG ACAACTATTC

2041 TAACACCTCA CCATTGAAAT TTATATGCCA CCTTGTCTTT CATAAAAGCT GAAAATTGTT
2101 ACCTAAAATG AAAATCAACT TCATGTTTTG AAGATAGTTA TAAATATTGT TCTTTGTTAC
2161 AATTTCGGGC ACCGCATATT AAAACGTAAC TTTATTGTTC CAATATGTAA CATGGAGGGC
2221 CAGGTCATAA ATAATGACAT TATAATGGGC TTTTGCACTG TTATTATTTT TCCTTTGGAA
2281 TGTGAAGGTC TGAATGAGGG TTTTGATTTT GAATGTTTCA ATGTTTTTGA GAAGCCTTGC
2341 TTACATTTTA TGGTGTAGTC ATTGGAAATG GAAAAATGGC ATTATATATA TTATATATAT
2401 AAATATATAT TATACATACT CTCCTTACTT TATTTCAGTT ACCATCCCCA TAGAATTTGA
2461 CAAGAATTGC TATGACTGAA AGGTTTTCGA GTCCTAATTA AAACTTTATT TATGGCAGTA
2521 TTCATAATTA GCCTGAAATG CATTCTGTAG GTAATCTCTG AGTTTCTGGA ATATTTTCTT
2581 AGACTTTTTG GATGTGCAGC AGCTTACATG TCTGAAGTTA CTTGAAGGCA TCACTTTTAA
2641 GAAAGCTTAC AGTTGGGCCC TGTACCATCC CAAGTCCTTT GTAGCTCCTC TTGAACATGT
2701 TTGCCATACT TTTAAAAGGG TAGTTGAATA AATAGCATCA CCATTCTTTG CTGTGGCACA
2761 GGTTATAAAC TTAAGTGGAG TTTACCGGCA GCATCAAATG TTTCAGCTTT AAAAAATAAA
2821 AGTAGGGTAC AAGTTTAATG TTTAGTTCTA GAAATTTTGT GCAATATGTT CATAACGATG
2881 GCTGTGGTTG CC**ACAAAGTG CCTCGTTTAC CTT**TAAATAC TGTTAATGTG TCATGCATGC
2941 AGATGGAAGG GGTGGAACTG TGCACTAAAG TGGGGGCTTT AACTGT**AGTA TTTGGCAGAG**
3001 **TTGCCT**TCTA CCTGCCAGTT CAAAAGTTCA ACCTGTTTTC ATATAGAATA TATATACTAA
3061 AAAATTTCAG TCTGTTAAAC AGCCTTACTC TGATTCAGCC TCTTCAGATA CTCTTGTGCT
3121 GTGCAGCAGT GGCTCTGTGT GTAAATGCTA TGCACTGAGG ATACACAAAA ATACCAATAT
3181 GATGTGTACA GGATAATGCC TCATCCCAAT CAGATGTCCA TTTGTTATTG TGTTTGTTAA
3241 CAACCCTTTA TCTCTTAGTG TTATAAACTC CACTTAAAAC TGATTAAAGT CTCATTCTTG
**3301** TCATTGTGTG GGTGTTTTAT TAAATGAGAG TTTATAATTC AAATTGCTTA AGTCCATTGA
3361 AGTTTTAATT AATGGGCAGC CAAATGTGAA TACAAAGTTT TCAGTTTTTT TTTTTCCTGC
3421 TGTCCTTCAA AGCCTACTGT TTAAAAAAAA AAAAAAAAAA AAACATGGCC TGAGAGTAGA
3481 GTATCTGTCT ACTCATGTTT AATTAAGGAA AAACACTTAT TTTTAGGGCT TTAGTCATCA
3541 CTTCATAAAT TGTATAAGCA CATTAAATAG CGTTCTAGTC CTGAAAAAGT CCAAGATTCT
3601 TAGAAAATTG TGCATATTTT TATTATGACA GATGTTTGAA GATAATTCCC CAGAATGGAT
3661 TTGATACTTT AGATTTCAAT TTTGTGGCTT TTGTCTATTA TTCTGTACTC TGCCATCAGC
3721 ATATGGAAAG CTTCATTTAC TCATCATGAC TTGTGCCATA TAAAAATTGA TATTTCGGAA
3781 TAGTCTAAAG GACTTTTTGT ACTTGAATTT AATCATGTTG TTTCTAATAT TCTTAAAAGC
3841 TTGAAGACTA AAGCATATCC TTTCAACAAA GCATAGTAAG GTAATAAGAA AGTGTAGTTT
3901 GTACAAGTGT TAAAAAAATA AAGTAGACAA TGTTACAGTG GGACTTATTA TTTCAAGTTT
3961 ACATTTTCTC CATGTAATTT TTTAAAAAGT AAATGAAAAA ATGTGCAATA ATGTAAAATA
4021 TGAAGTGTAT GTGTACACAC ATTTTATTTT TCGGTATCTT GGGTATACGT ATGGTTGAAA
4081 ACTATACTGG AGTCTAAAAG TATTCTAATT TATAAGAAGA CATTTTGGTG ATGTTTGAAA
4141 AATAGAAATG TGCTAGTTTT GTTTTTATAT CATGTCCTTT GTACGTTGTA ATATGAGCTG
4201 GCTTGGTTCA GTAAATGCCA TCACCATTTC CATTGAGAAT TTAAAACTCA CCAGTGTTTA
4261 ATATGCAGGC TTCCAAAGGC TTATGAAAAA AATCAAGACC CTTAAATCTA GTTAATTTGC
4321 TGCTAACATG AAACTCTTTG GTTCTTTTAT TTTTGCCAGA TAATTAGACA CACATCTAAA
4381 GCTTAGTCTT AAATGGCTTA AGTGTAGCTA TTGATTAGTG CTGTTGCTAG TTCAGAAAGA
4441 AATGTTTGTG AATGGAAACA AGAATATTCA GTCCAAACTG TTGTAAGGAC AGTACCTGAA
4501 AACCAGGAAA CAGGATAATG GAAAAAGTCT TTTAAAGATG AAATGTTGGA GCCAACTTTC
4561 TTATAGAATT AATTGTATGT GGCTATAGAA AGCCTAATGA TTGTTGCTTA TTTTTGAGAG
4621 CATATTATTC TTTTATGACC ATAATCTTGC TGTTTTTCCA TCTTCCAAAA GATCTTCCTT
4681 CTAATATGTA TATCAGAATG TGGGTAGCCA GTCAGACAAA TTCATATTGG TTGGTAGCTT
4741 TAAAAAGTTT GTAATGTGAA GACAGGAAAG GACAAAATAG TTTGCTTTGG TGGTAGTACT
4801 CTGGTTGTTA AGCTAGGTAT TTTGAGACTA CTTCCCCATC ACAACAACAA TAAAATAATC
4861 ACTCATAATC CTATCACCTG GAGACATAGC CATCGTTAAT ATGTTAGTGA CTATACAATC
4921 ATGTTTTCTT CTGTATATCC ATGTATATTC TTTAAAAATG AAATTTATAC TGTACCTGAT
4981 CTCAAAGCTT TTTAGCTTAG TATATCTGTC ATGAATTTGT AGGATGTTCC ATTGCATCAG
5041 AAAACGGACA GTGATTTGAT TACTTTCTAA TGCCACAGAT GCAGATTACA TGTAGTTATT
5101 GAGAATCCTT TCGAATTCAG TGGCTTAATC ATGAATGTCT AAATATTGTT GACATTAGGA
5161 TGATACATGT AAATTAAAGT TACATTTGTT TAGCATAGAC AAGCTTAACA TTGTAGATGT
5221 TTCTCTTCAA AAATCATCTT AAACATTTGC ATTTGGAATT GTGTTAAATA GAATGTGTGA
5281 AACACTGTAT TAGTAAACTT CATCACCTTT CTACTTCCTT ATAGTTTGAA CTTTTCAGTT
5341 TTTGTAGTTC CCAAACAGTT GCTCAATTTA GAGCAAATTA ATTTAACACC TGCCAAAAAA
5401 AGGCTGCTGT TGGCTTATCA GTTGTCTTTA AATTCAAATG CTCATGTGAC TTTTATCACA
5461 TCAAAAAATA TTTCATTAAT GATTCACCTT TAGCTCTGAA AATTACCGCG TTTAGTAATT
5521 ATAGTGGGCT TATAAAAACA TGCAACTCTT TTTGATAGTT ATTTGAGAAT TTTGGTGAAA
5581 AATATTTAGC TGAGGGCAGT ATAGAACTTA TAAACCAATA TATTGATATT TTTAAAACAT
5641 TTTTACATAT AAGTAAACTG CCATCTTTGA GCATAACTAC ATTTAAAAAT AAAGCTGCAT

5701 ATTTTTAAAT CAAGTGTTTA ACAAGAATTT ATATTTTTTA TTTTTTAAAA TTAAAAATAA
5761 TTTATATTTC CTCTGTTGCA TGAGGATTCT CATCTGTGCT TATAATGGTT AGAGATTTTA
5821 TTTGTGTGGA ATGAAGTGAG GCTTGTAGTC ATGGTTCTAG TGTTTCAGTT TGCCAAGTCT
5881 GTTTACTGCA GTGAAATTCA TCAAATGTTT CAGTGTGGTT TTCTGTAGCC TATCATTTAC
5941 TGGCTATTTT TTTATGTACA CCTTTAGGAT TTTCTGCCTA CTCTATCCAG TTGTCCAAAT
6001 GATATCCTAC ATTTTACAAA TGCCCTTTCA GTTTCTATTT TCTTTTTCCA TTAAATTGCC
6061 CTCATGTCCT AATGTGCAGT TTGTAAGTGT GTGTGTGTGT GTCTGTGTGT GTGTGAATTT
6121 GATTTTCAAG AGTGCTAGAC TTCCAATTTG AGAGATTAAA TAATTTAATT CAGGCAAACA
6181 TTTTTCATTG GAATTTCACA GTTCATTGTA ATGAAAATGT TAATCCTGGA TGACCTTTGA
6241 CATACAGTAA TGAATCTTGG ATATTAATGA ATTTGTTAGT AGCATCTTGA TGTGTGTTTT
6301 AATGAGTTAT TTTCAAAGTT GTGCATTAAA CCAAAGTTGG CATACTGGAA GTGTTTATAT
6361 CAAGTTCCAT TTGGCTACTG ATGGACAAAA AATAGAAATG CCTTCCTATG GAGAGTATTT
6421 TTCCTTTAAA AAATTAAAAA GGTTAATTAT TTTGACTA

**e.** *RARB* 3'UTR sequence with underlined bioinformatically predicted hsa-miR-548ba target sites. microT CDS v5.0 predicted target sequences locations are marked in green: positions 816-835 and 1257-1270. TargetScan and miRDB predicted seed sequence location is marked in red: position 1264-1270.

1 GACATTTTCT AGCTACTTCA AACATTCCCC AGTACCTTCA GTTCCAGGAT TTAAAATGCA
61 AGAAAAAACA TTTTTACTGC TGCTTAGTTT TTGGACTGAA AAGATATTAA AACTCAAGAA
121 GGACCAAGAA GTTTTCATAT GTATCAATAT ATATACTCCT CACTGTGTAA CTTACCTAGA
181 AATACAAACT TTTCCAATTT TAAAAAATCA GCCATTTCAT GCAACCAGAA ACTAGTTAAA
241 AGCTTCTATT TTCCTCTTTG AACACTCAAG ATTGCATGGC AAAGACCCAG TCAAAATGAT
301 TTACCCCTGG TTAAGTTTCT GAAGACTTTG TACATACAGA AGTATGGCTC TGTTCTTTCT
361 ATACTGTATG TTTGGTGCTT TCCTTTTGTC TTGCATACTC AAAATAACCA TGACACCAAG
421 GTTATGAAAT AGACTACTGT ACACGTCTAC CTAGGTTCAA AAAGATAACT GTCTTGCTTT
481 CATGGAATAG TCAAGACATC AAGGTAAGGA AACAGGACTA TTGACAGGAC TATTGTACAG
541 TATGACAAGA TAAGGCTGAA GATATTCTAC TTTAGTTAGT ATGGAAGCTT GTCTTTGCTC
601 TTTCTGATGC TCTCAAACTG CATCTTTTAT TTCATGTTGC CCAGTAAAAG TATACAAATT
661 CCCTGCACTA GCAGAAGAGA ATTCTGTATC AGTGTAACTG CCAGTTCAGT TAATCAAATG
721 TCATTTGTTC AATTGTTAAT GTCACTTTAA ATTAAAAGTG GTTTATTACT TGTTTAATGA
781 CATAACTACA CAGTTAGTTA AAAAAAATTT TTTTAC**AGTA ATGATAGCCT CCAAG**GCAGA
841 AACACTTTTC AGTGTTAAGT TTTTGTTTAC TTGTTCACAA GCCATTAGGG AAATTTCATG
901 GGATAATTAG CAGGCTGGTC TACCACCTGG ACCATGTAAC TCTAGTGTCC TTCCTGATTC
961 ATGCCTGATA TTGGGATTTT TTTTTCCAGC CTTCTTGATG CCAAGGGGCT AATTAATATT
1021 AACAACTCCC AAAGAAACAG GCATAGAATC TGCCTCCTTT GACCTTGTTC AATCACTATG
1081 AAGCAGAGTG AAAGCTGTGG TAGAGTGGTT AACAGATACA AGTGTCAGTT TCTTAGTTCT
1141 CATTTAAGCA CTAGTGGAAT TTTTTTTTTT TGATATATTA GCAAGTCTGT GATGTACTTT
1201 CACTGGCTCT GTTTGTACAT TGAGATTGTT TGTTTAACAA TGCTTTCTAT GTTCATA**TAC**
1261 **TGTTTACCTT** TTTCCATGGA GTCTCCTGGC AAAGAATAAA ATATATTTAT TTTAAAAAAA
1321 AAAAAAAAAA AAAAAAAAAA AAAAAAAAAA AAAAAAAAAA AAAAAAAAAA AAAAAAAAAA
1381 AAAAAAAAAA AAAAAAAAAA AAAAAA

**f.** *SP110* 3'UTR sequence with hsa-miR-548ba target site at position 33-40 predicted by microT CDS v5.0 and marked in green.

1 ATTTCTACTA CCCTCTCAGT CACCATGTTG CA**GACTTTCC** CTGTCTGGAG GCTCACCTTA
61 GAGCTTCTGA GTTTCCAAGC TCTGAGTCAC CTCCACATTT GGGCATGGCA TCTTCAAAAC
121 AATTAATTTG CATAGTTAAT TTGGGATGGG GAAGCAAATG ACTCTAAAAT AAAAATTAAA
181 TGAAAAAGCT CAAAAAAAAA AAAAAAAAAA A

**g.** *ADAM19* 3'UTR sequence with hsa-miR-7973 target sequence commonly predicted by TargetScan and miRDB is marked in red at position 624-630.

1 ACCTGTCCAA GGGGCTTCTC CCTTTCCTTG AGCTCTCTGG ACACTGCAGA GGACCCATGG
61 CCATGGAACC CTGAAGAAGC ATGTCTGGCC GCCTCTGAGC TCCTCCCACC CTCCTCCAGG
121 AACCTCCACA TCTCCAAAAA TCTCCCTGTT GACTCAGTGC CTCCTCGGCT TCCTTGGAAG
181 CCCAGAGGGA CTATGATCTG ATGGCCTCTA GGTGTTGTTT TGTGCAATAT ACAGCCCCAG
241 GTAGGGAGGG GAGAGTATGA GGAGGGTGAC TGGCAGCTTC TCCTCCAGAC TCCTAGCCCC
301 GAGGTGCTGA TGGAGATGCT CAAGGCCAGC AAGCCCCTCA GGCCAGCACT TCGCTTGCAG
361 AAGCCATCCA TTCACTCCTG GGGTGCAGGG CACGCAAGAG AGCTTCCCAT TGCTTCTGCT
421 CTCCTCAGAG GTCCCGGGCT GGATGGAGGC TGGTACTTAC CCACCCCTTT TAGCTTTTAG
481 GGATTAAGGA AGGGTCAAGC CAGCCACTGC TGTGGCCCTG CCCAGGGCTT GGTTGAGGGA
541 ACGGCTTCTG GCTGTATGGC TGCATGTGAC AAGCCACGTC CCCTCCCACC TCTCCCCAAA
601 CCCCTGCATC CCTGTATTCA CAC**GGGTCAC** TCTGACTCAG ACAGGTACTA TTCGTAGGCA
661 GTGTAGACAG CAGGAGGAGC ACCGGGCTTG GGCTTCCTCT GAGCCGTGAT GCCAAAGGTT
721 GCGACTCCTG ACTCTGGATA ATTTTTAGTT GCTCTTTGTT TTCTCTGCCG CACTTTCCTG
781 GTGCCCCACG CTTTTCTCTC TTCCTTCCCC TCTCATTCTC CCTCTAATGT GTGGTGCTTT
841 GGTGAGCAAA CCCTCAGCAG TCCTGACCTT CGGGTGACCA GGTGCTTGTG ACCTACAAGT
901 CAGAGTCCTC TCTCACAGTC GGCCACTGGA TTTCCCTCAC TGGCTCTCAG GAGTGTGACC
961 AGAGTAGACT TGGGGCATGG CCATTGGGGT CATATGTTTA TTTTTCATTG TGTTTTGTGA
1021 CCTCAGCAGG GTGGGGGTCT TCCTCCTTAC TCTAAGCTAA ATCTAGGTGA GGTTTCCCCT
1081 TAGGGAGCCC AGCTATTTAC AAAGTACACA CGAGGGAGCA GGCTGGTCAT TGACTTCGGG
1141 CTGGACCGTT GCCCTCTGAG CAGAGAACAG ACCCATTTCT GGGAGCTGCC CGAGATCACT
1201 GGAGAAGGCA GCCAGCAGCA GCTGCACTGG AACAGTCAGA GCAGGGAGCC TCTTCCTCAA
1261 CCCAGCTTTT TGTCATTCAC TTCCTTTTGT TCTCTCTCTG GTCACTGCCC TTACCTGACC
1321 CTCACAGAAA GAGAGCTCTG AGCAGGTGAG GGGGTCTGCG GTGGCTCCTG TCTTCCCTGC
1381 AGCAGGGAAG GAGGGCCGTG TGGTGCTTTG CTAGATAGGA CGGTTTTTGC AAAGCACCTG
1441 GAGATGTTTG CTGGGAGATA GACTCCCACT CCACAAAGGT GCTGGGTGGC TCTCCGGACA
1501 GGAGCTGGCC TGACTCTCAC TCCTCTGAGG CTTTCCTGGG GCCTCCTCCC ATCCTGCCAT
1561 GAGCAATTGT TTGCTCTTGA AAACCTCACT GCAAGGCTGA GGCTGAGCTT CTGATTCACC
1621 ACCCCAGGGC CTCCTTATAG TTCTCTGCAC ACAATAGGTG CTTCTTGGAT GTTCTTGGGT
1681 TTGGAAATAA GTGGAAAATA CGGGATGTAC CCCTGGGGGA AAAGCCTGGG TTGGGTTTAG
1741 AAAGATCTCA GGAAAATGAG TTTCTCTTCC CTCAGGGTGG CTGTGATACA GGTTCCCCAT
1801 GTCCTTGCCG TGGGTCATCC TTGCTGTGGG TCATCCTTGC TGTGGAGATC CATTCCCCAC
1861 CTTTCCTGTG GCCCAACCTT TTATTTAAAT GTGCTACCCT CTGCCTCAAG GCTTGGTTCC
1921 TGGAAAGTAA AGGTGAAAAC ATCCCCTTTC ACCCCTCTGC AAAACAAACA AGCAACATCC
1981 TCAAAACCCA ACCCCATGCC TCACAGAGCT TCCTGTGGCT TCTCCAGCCT TTCTCCCTCA
2041 CATCAGGAGG TAGATAGCTC TGAAATGACA GCGCCACAGC CATAGTGACT GCATGAGCCA
2101 TCTGAACCTG CAGTCCACCC TCCCTGGAAC CACACCAGAA AGAGACCTGG GTTGTCGTTT
2161 TCTTGCTTTT TGTTTTGTTT TGTTTTATTA TTTTCATATC ACCTCCATCC CATAAAGTTG
2221 TACTGTGAAC TGGAAGATGG TGGAATGTTT TGGAATTTGA TAGACTTTCG GCAACCAGTT
2281 CTACTAATGC TTCACTCCTG GCTCTGTTCA GGGAGGCTGC CCAGGAGGAA GACTGGCCAT
2341 TATGCATCCC CTTTTCTTTC CAGTGCCCAG TATGCTGTTT TGAGGTGTCA AATACAAATA
2401 AATCTGGGCT TAGGGAAGGA GAGACCTTAT TCCAAAGCAC GATTGCAGAA GGGGAAAGGG
2461 AATATTGCAA AAGGGAGAGG AAGGGGCCTT ATGGGAATAG TGAAAAGGCT CAGACCGACC
2521 GATGGCAAGA TCTGCAAGCG TCTCAAAGCC CAGGCAGAAA AGGACTTTTC TTTTATTGGA
2581 AGAAGTAAAC ATGGCTAGAA AGAACCACGT TCAGGGAATG ACGTTGTGCC CAGCCTTTTT
2641 TTTTTTTTTT TTTTTTTTGT CTCCAGGGGA GGGGCTGTTT GCTGGCTCAG GCTGAGGATG
2701 GCCCAAAGTC CAGGGTCTGG TGGGGAGGAG GGAAGCTTAA CTCAAGTTTG GGTTAGTGAG
2761 TTAGCAAGCT CTTTGTGCAG ATGGGGATGT AGGTAAATCT TTTTAAAAGT GAAATTAACC
2821 TCCTGCCAAT TTTACAACCC AAGAATTTTT TTTTAAGGGC CTTGGAGCCA TCTCTAAAAC
2881 AAACCTCAAG GGATTTAGTG CCCTGTCTCC CTGTCTCTAG AAGCCTTAGC CTGGGCACCT
2941 GGCTCAATCT TGTAACTGCC TGCTAGCCAT AGATTCCTTT CAGCCTTGCT GACTTCTCCC
3001 TATAAAAGTA AAGCCTTTTT CTGCCCCAGC TCTGAGACAC TTGCAGATCT TAAGGTCTGA
3061 GACTTGCTGA TTTTCTGGTT GGAGTGTTTT TTTGTATTGC CATAGTCCCT TCCCCCTGAA
3121 GCAATAGCCC CTCCCCACCT CCTGCAATAC GCCTTTCCAA TCTTTATTGG AAGTCTCTCC
3181 CTGCCTACTT CCTAATTTAT TCTTATTTGA CAGAGGGTAT GGAAGACTTG CAATTTGAAA
3241 ACTGGGGACC AGTTCCAAAG TCAGTAATTG TGTTAACCAC GTGTATAACA GCTCTGCTGG
3301 ACACCCAAGA AAGCCATGGG AACGCCAACT GGAAAGGTCC CCTTCCCCAG GGGAGCCTGC
3361 GAAGGAGAGG TTCTGTAGAA TCCAAGCCCA CATTTCCAAA GTCACCCCCA ACGCGTCCTC

3421 TCACACCGTC CACTGTGCGT TTGTATGTGT CTGGGATCCA GGGCAATGTG AATTTTCTTT
3481 TTATTTGGGA GATTGTTCAC GGAAAACAGA TCTTCTTCTC TCTTGTCCAC CTATTAATTG
3541 TTTACAATAT TTGTACATCT ATGCAAAATA CTTGAATGGG CCATGGTGCC TTTTTTCCTT
3601 GTTAGTATTT AATTAAAAAT GAATTGTTTG TCATTTGCAA TGTTAAAAAA AAAAAAAAAA
3661 AAAAAAAAAA

**h.** *ATHL1* 3’UTR sequence with hsa-miR-7973 target sequence predicted by TargetScan is marked in blue at position 346-352.

1 TCAGGAACGG TGGCTTCAGA GACGTCTCTT GGGCCTTCCC TCTGGCCACG TCTGCACCCA
61 CCCCTCCTGG GCACCCTCCT AGCCTGCCAT CCCTCACCTG CAGCCAGGCT CTCAGGGAAG
121 GTCCATGCTG CTTGGCCTGA GTTCAAGGCT TTCTGCCTGT AGCCTGGACT CCCGTGGACC
181 CCCGTGGGCA GGTGGCTTCC CCGTGGCATC TCCACACCGC CTCTGCCTGC CCCTGTGGAC
241 TGATGCTATC GCGCACCGTC CCACGACCCC ACCCCGAGCT CCTGAAGCCG GGGTCTGAGC
301 CTGCATCACC TCTGGCCTCT CATCCCCCAC TCTCCTGAGA GCAGT**GGTCA CA**GCGGCCGG
361 CCGCTCTGCT GAGAAGGCAG AGAGGCAGGC TCAGGCCTCA GCGTGGACAG CAGGGATAAG
421 GGGCACGAAG GACGGGGACT CGGCCCCTTC AGAATTCCTC AGGACTCTCA GGTGCAGCTT
481 TGCCAAAAAG GAACTTTTCA TGTCATGCAG TTGAGGGGAC TTAGTCTCAA TCCCAGGCTC
541 CTCTTGACTC TGGGCAGCTT TAATCAGGTT GGGCAGCCTC TGCTACAGCG TGGGGTGGGA
601 TGGCTCTCTT CCCTCAGCCA CGCCGCTTGT GAGGACAGAG GTGGGGGAGT GGGAAGTGGG
661 AAGTCACCAG AGAACAGGAG AGGGATTTGA GGGCGAGACC CCAGCGCTCT CCACGGACCA
721 GCCAGAGGGA CTGGAGCCAG GTGTGCATGG GTTCAAGGCC CTGGCCCTGC CCAGCCTTTG
781 TCTTGGGAGC TCAGCCCCAG GGTTCGGTCG TCAGCAGTTT CCCAAGAACA AGATGTGATG
841 GCATCTGCTG CTGAAACCCT GATGAGGACC AGGCCCCCTG CACCGCTGTC AGCCTGAGGA
901 ATTAAAGCTT TGGTGCTGGG GAGAGCATTA TTCCTCTGAA A

**i.** *ATP6V1A* 3’UTR sequence with underlined bioinformatically predicted hsa-miR-7973 target sequences. microT CDS v5.0 predicted target sequence locations are marked in green: positions 1336-1352, 1543-1553 and 2307-2323. TargetScan and miRDB commonly predicted target seed sequence locations are marked in red: position 1547-1554.

1 AAGCCTTGAA GATTACAACT GTGATTTCCT TTTCCTCAGC AAGCTCCTAT GTGTATATTT
61 TCCTGAATTT CTCATCTCAA ACCCTTTGCT TCTTTATTGT GCAGCTTTGA GACTAGTGCC
121 TATGTGTGTT ATTTGTTTCC CTGTTTTTTT GGTAGGTCTT ATATAAAACA AACATTCCTT
181 TGTTCTAGTG TTGTGAAGGG CCTCCCTCTT CCTTTATCTG AAGTGGTGAA TATAGTAAAT
241 ATACATTCTG GTTACACTAC TGTAAACTTG TATGTAGGGT GATGACCCTC TTTGTCCTAG
301 GTGTACCCTT TCCTCATCTC TATTAAATTG TAAACAGGAC TACTGCATGT ACTCTCTTTG
361 CAGTGAATTT GGAATGGAAG GCCAGGTTTC TATAACTTTT GAACAGGTAC TTTGTGAAAT
421 GACTCAATTT CTATTGTGGT AAGCTCATTG GCAGCTTAGC ATTTTGCAAA GGAATTGCTT
481 TGCAGGAAAT ATTTAATTTT CAAAAACATA ATGATTAATG TTCCAATTAT GCATCACTTC
541 CCCCAGTATA AATCAGGAAT GTTTGTGAGA AACCATTGGG AACTATACTC TTTTTATTTT
601 TATTTTTTAT TTTTTTTATT ATTTTTTTTT TGGGGACGGA GTGTCCCTCT TGTTGCCCAG
661 GCTGGAGTGC AATGGCGTGA TCTTGGCTCA CTGCAGCCTT CGCCTCCCGG GTTCAAGTGA
721 TTCTCCTGCC TCAGCCTCCC GAGTAGCTGG GATTACAGGC ATGCTCCACC ATGCCCAGCT
781 AATTTTGTAT TTTTAGTAGA AACGGGGTTT CACCATATTG GTCAGGCTGG TCTCGAACTC
841 CAGACCTCAG GTGATCCGCC CACCTCGGCC TCCCAAACTG CTGGGATTAC AGGCGTGAGC
901 CACCGCGCCT GGCCAGGGAC TATACTCTTT TTAAAATAGA CATTTGTGGG GCTCACACAA
961 TATATGAAAT AGTACCCTCT AAAAAAGAGA AAAAAAAAAT CAGGCGGTCA AACTTAGAGC
1021 AACATTGTCT TATTAAAGCA TAGTTTATTT CACTAGAAAA AATTTAATAT CAAGGACTAT
1081 TACATACTTC ATTACTAGGA AGTTCTTTTT AAAATGACAC TTAAAACAAT CACTGAAAAC
1141 TTGATCCACA TCACACCCTG TTTATTTTCC TTAAACATCT TGGAAGCCTA AGCTTCTGAG
1201 AATCATGTGG CAAGTGTGAT GGGCAGTAAA ATACCAGAGA AGATGTTTAG TAGCAATTAA
1261 AGGCTGTTTG CACCTTTAAG GACCAGCTGG GCTGTAGTGA TTCCTGGGGC CAGAGTGGCA
1321 TTATGTTTTT ACAAA**ATAAT GACATATGTC AC**ATGTTTGC ATGTTTGTTT GCTTGTTGAA
1381 TTTTTGAACA GCCAGTTGAC CAATCATAGA AAGTATTACT TTCTTTCATA TGGTTTTTGG
1441 TTCACTGGCT TAAGAGGTTT CTCAGAATAT CTATGGCCAC AGCAGCATAC CAGTTTCCAT

```
1501 CCTAATAGGA ATGAAATTAA TTTTGTATCT ACTGATAACA GAATCTGGGT CACATGAAAA
1561 AAAATCATTT TATCCGTCTT TTAAGTATAT GTTTAAAATA ATAATTTATG TGTCTGCATA
1621 TTGCAGAACA GCTCTGAGAG CAACAGTTTC CCATTAACTC TTTCTGACCA ATAGTGCTGG
1681 CACCGTTGCT TCCTCTTTGG GAAGAGGAAA GGGTGTGTGA ACATGGCTAA CAATCTTCAA
1741 ATACCCAAAT TGTGATAGCA TAAATAAAGT ATTTATTTTA TGCCTCAGTA TATTATTATT
1801 TAATTTTTTA GGTAATGCCT ATCTCTTGGT CTATTAAGGA AAGAAGCAAT CAGTAGAGAA
1861 TTCAGGATAG TTTTGTTTAA ATTCTTGCAG ATTACATGTT TTTACAGTGG CCTGCTATTG
1921 AGGAAAGGTA TTCTTCTATA CAACTTGTTT TAACCTTTGA GAACATTGAC AGAAATTATG
1981 CAATGGTTTG TTGAGATACG GACTTGATGG TGCTGTTTAA TCAGTTTGCT TCCAAAGTGG
2041 CCTACTCAAG AGGCCCTAAG ACTGGTAGAA ATTAAAAGGA TTTCAAAAAC TTTCTATTCC
2101 TTTCTTAAAC CTACCAGCAA ACTAGGATTG TGATAGCAAT GAATGGTATG ATGAAGAAAG
2161 TTTGACCAAA TTTGTTTTTT TGTTGTTGTT GTTGTTTTGA ATTTGAAATC ATTCTTATTC
2221 CCTTTAAGAA TGTTTATGTA TGAGTGTGAA GATGCTAGCG AACCTATGCT CAGATATTCA
2281 TCGTAAGTCT CCCTTCACCT GTTACAGAGT TTCAGATCGG TCACTGATAG TATGTATTTC
2341 TTTAGTAAGA ATGTGTTAAA ATTACAATGA TCTTTTAAAA AGATGATGCA GTTCTGTATT
2401 TATTGTGCTG TGTCTGGTCC TAAGTGGAGC CAATTAAACA AGTTTCATAT GTATTTTTCC
2461 AGTGTTGAAT CTCACACACT GTACTTTGAA AATTTCCTTC CATCCTGAAT AACGAATAGA
2521 AGAGGCCATA TATATTGCCT CCTTATCCTT GAGATTTCAC TACCTTTATG TTAAAAGTTG
2581 TGTATAATTG TTAAAATCTG TGAAAGAATA AAAAGTGGAT TTAAATTAAC AAAAAAAAAA
2641 AAAAA
```

**j.** *FMNL3* 3'UTR sequence with bioinformatically predicted hsa-miR-7973 target sequences. microT CDS v5.0 predicted target sequence locations are marked in green: positions 805-824, 2481-2493, 3760-3772, 3894-3903, 4262-4269, 4657-4682, 6248-6260, 7301-7314 and 7635-7642. TargetScan predicted seed sequence locations are marked in blue: positions 233-239, 1620-1626, 3606-3612, 3974-3980, 5387-5393 and 7063-7069.

```
   1 GACCCTCTCG GAGGCAGAAG CACTTCACCC CTCAGAGTCC TACAAGTCCA ACCAGTGGAC
  61 CTGGAATTGG CCAAGGGCTC AGGAGAGGGC TGTGTTGCTC TCTCAACCAT GTCCGCCCCA
 121 GCTCTTGAGG CTGGATCTTT CTACTTGTGC CACTATGGGC ACTAGGTCTG TAGGTCCCTG
 181 GTGCTTCCAG TACTGCACTT TGCCCCCAAG GTCTTTGACT TCATTTCCTG AAGGGTCACC
 241 AAGGGTCAGC TCAGGGCTAG GAAAAACCAT TTCCAGCCTT AGCCTAAACA GGGCACATAG
 301 ATCTCTCCCA CTAGAGCCCA GAGGGATGAG AGGAGAGAGC AACTGTTCTT TCTTTTTTTT
 361 TCCCTTCCTT TTTGCAGGCC CATGGTCAGC TCTGGTCTAG ATGTATCTAG GCTTAGGGAG
 421 CCCTGGGACC CTGCCAGTCT CAGTTCATCT CTTACCAGGA GCAAGGCCCT CTGAAGCATG
 481 GGCCGTGCCA GCTGTGCCCT AGTGATAAGG GTAAGAGGAG AGATGTCAGG CTCACTGCCC
 541 TTATTTCTCT GCCCATTTCA TTCTAGGCTC ACACTGTCCT TGTCAAGGTG GGCAGAAGAG
 601 CAGAGGTCTT CTCTGCACCA ACCACTCTCT GCAGATGGCT GGAGAAAGGG TGTGCCAACT
 661 CTTCACCTTC TTTCCTCAGC TACGGTTTTT TTTTTTTTTT TTTTTTTTTT GGCAGGGGAC
 721 AAGGAGCATG GTGGTCTTGG CTATTTGCTT ACCTTCCCGT TTCTCCTCTG CCCCTGGAAG
 781 GGAATGTGGG GGCCCACTTT TTTGTACATG TACCACCTCC CTTTCCTCTT ACTGTACATA
 841 AACCTCAGAC TCTCCCCCTC TCAACAAAGG CTTGATGCAC CAGGCCTGAC TTTGTCTCCT
 901 CTTTCTGAGC TTAGGGGGCT AGGGTGGCCA TTTAGTCTGA CCTGGGCTAT GGGGATAGAA
 961 AAGAAATCTT TGGGGCACTG CATCTGTTTT GGAGGAGATG AGATCAGAAC TCTGGGGAAA
1021 GGAGAACCGG CTAGGTGGGC TCTAGCCACA CAAAGGGAAG CAGGCCTGCC ATGAGACACT
1081 AGTGCCCTCT GCTGGGCATG TGTCCAGTCC CCAGCCTGCC CCAGTAGCAC ATGGAAAAGA
1141 GTCACTGGCC ATGTTATAAA TATTGTTATT TAAAAAACAA AAAACAAAAC ACACGTACAT
1201 TAGGTCCTGG GTGGAGGAAG AAGTGGTGGA GCCTCAGGGC CAGGGCAGGC AGGAGCAGGC
1261 ACTGATGCTA GAGGGTGATG ACCCTGCCCT CTGCCACCCA GGCTGCGTCC AACAGCCTGT
1321 GACTCTTCCC TGGAGACCCC TTCTCTCACA CTTTTATCTA CTGGCATCAC CCACCCCACA
1381 TGGCATATAC CACCCATCTC TGGTCTGGGG CTGAGGGCAA GGACTGCACT AGTCCAGGGA
1441 GTAGGGGACC CTTGAACACC AAGAGCAACC TTGAGAGGTG CCAACAATGG TGTGCTGGGG
1501 GTGGGGTTAC TTTAGAAAAA GTGGACACAG ACCAGACTAA GGAAGAAGTC TGAGGCAGAC
```

1561 GGTGAGGGTG GCCCAGGAGC CATGGCCTCA CACAGACCCG AGGCAGAGAA CAGCTCATT**G**
1621 **GGTCAC**TGGT GATCATCCAG CTGCTGTAGG AGTGTCCGCC GCCGCCTCTC CAGCTCACCC
1681 TCACTCAGCT CACTTTCTGA CGTGTCCCAG CCTGTCTGTT GGAGCAAAGG CACAGTTCAC
1741 CCTTAGCCTG CACTAAGGGC CGTGGAAGCT TGACCAGCGT GCTGACTGGC CTAGGGCCCC
1801 TGCCCCTCAC CTTCTCCTTC TTGATTCCAA AGCCTGGGGA ACGGTTAGGG AGCTCTGCCT
1861 GTTGGAGCTC CCTGTCCTTG TCCTGTTCTT GTTCTTTCTC ATCGCTCTCC TTGCCAGCTT
1921 TCTCCTCAGG GTCTGTCTCA CTCTCAGGAC TATTCTGTAA GAGTCCAGAG CCCACCTCCT
1981 CAGTTCCTGT GGATAGCCTG GGAGGCAGGT AGAACAGAAC TCACAAAATA GCAGGGGCTT
2041 AGGTAGAAGT TCCTTCACTC ACCGACTTGT GTCTTCTCTT CTTAGTTTTC TTTTTTGGTT
2101 TCTTGGCTTT CCGAAGGCCA TGATCTAGAG ATAGAAAAAT CCCTACTAAG TCCCAACAGG
2161 AGAAAGACAG GCTCTTTTTC TTTGTTCTAG CCCCTCTGTT CATTCTTCAC AGGCAGGAAC
2221 CTCATGCTCT GAGGAAGTCT AAGTCCAGTT TTCCTTCTCC AGCTTCTTGC TTGGGCAGGT
2281 AGAGGAAGTT CTTTCTACTT CCACAGGTGC AGCCCTCCCA ATCCAGTCTA CCCTCAAAGG
2341 AGCTGGGGCT TGTGGAGGGC ATCCTCTCAG TCTGGGACAA CAGTTCTCCA GCTTCTACGC
2401 TCACAGCAAC TGCTTACCTG CTCCAAGAAG ATGGGAGGAA GGGGAGCCCC GTCCTCCAAG
2461 GGCAGCACCC CCACTTTCAA **CTGAATCAAG TGA**GGAAGAG GGCTCAGAGC CTGACTCTGA
2521 GGGGTTCCGC CTCCTCCGCT TGGGGGGCCG GAGAGATGGT GGGGGCAGCT CCTCTTCTTC
2581 TGACTCAGAG CCCTGGGCAG ATCAGCAGAT ACATGTCAGT GCAGAGGTTC TCTGCCCTCC
2641 AGAAGCCCTC GTCTGGGAAT TCGAGAGAGG AGGAGGACCC AGGGCCAAAG GAATCCTCAA
2701 ACTTCCCTCC AAGGGAGATA TTGTAAGGGG AAAACAAAGT GTTTATTTCC CCAAAACAAC
2761 CCAGCTTCAC CCCATGTCCC TTCTACACGC TTACTCACTG AGGGTGAGTG GGAACGCTTG
2821 TGATGGTGCT TCTTGCCTTT CCTGCCATGC TTTCGGCCTT TGGTGTGGAG GTGCTGGCAT
2881 TCAGTCTGCT AGAGGCAGAG GAAGAAGGGT GAGGACAGCA CCAGACTTAG CCCCAGGAAG
2941 GATCCTACTT CCTAAAATCT GGCTTTGCCT CCCTTTCTCA GACTAGAGCC TTCTGCCTAT
3001 CCCAGGGATC CTCAGGCCTC CCCTTCCTTA GCTGGCCAGG AGTGCCCTAA GGGTAAGAGG
3061 GAGGCCACTG AGGTCTCTGT TGAACTGGTC TAATGAGTTC AAGGGCCTGA GGCAGATCCA
3121 GAGGACAAGC CTGCCTCACC TCCAGCACCT GTAGGAACTC CCGGAAGAGC CGGATCCGCT
3181 CCGACTCCAG GGTGATCTGC TCAAAGGCTG AGTCACACAC AAAACGCTCA CGGACCTTGA
3241 ACGAGGAGAG CACTGCTGAT CAGACTAGGC CCCACTCAGC CTGGCCTGGA CACCGCCATG
3301 AGCACAGGGC AGGGCTGGGG CAGAGAAAGG AAGGGAACTG GAGCTGGCCC AGGGCTGTGA
3361 CGTGACTGGG GTGGGCCCTC AGTGGTGAAG CACCGCTGCC CTCAGGCTTG AGGGGTGCTT
3421 GGGGCCAGGC TACGCTCCTG ACCTCTTCCC AGGCAGTGCC TAGCTCCAGA GCAGGCACAG
3481 CCTGCCTCAG CATGCTTCGA AAGGCAGCTT CCCTGCGCCG CATCCTGCGT GCCTCCTCCT
3541 TCTCCCGCTC CCTCTCCCGT GCCTCTGCTT TCTCCAGCAG CTGAGGAAAG AAGGGACACG
3601 GAACA**GGGTC AC**CCAGGAGG GTGGATGCCA GGTCCCACTC AGGCATACCT TAGGAGGTGG
3661 AAAGTTAGGG TCTGGGCTTC TACCACCATC CCCTGAATTG GGAATATAGT CAGTGGGACA
3721 GTGAGTGAGA AGGAATGGAG CAAGACACAG ATGAATTAGA **GAACTTCCCA CG**CCCCGCCC
3781 AGCCCCTCAC ACTATTGAAG GTCAGCTTGA TGTTGCCTGC GTCCAGTGCG GCAGCCCTCT
3841 TGTCAAAGCT TATGACGTGG GCGAAGTCCT CAAAGGCCGT GTTCACCTCC ACG**CAGAAGC**
3901 **CCC**GGTCCTG TGGGTACAGC AGTGCATGAA GCAGGGGCAC CACGTGCCTG AGGCCAAAGC
3961 TGGCAGGCAG AGT**GGTCACA** TTACAGATGG TCAAGAAGGC CCTTGCCTCT GGAAGAGATC
4021 CACAGGACTG TGGTCTGCAT TAGAAACCAA GTTATCCAAG TCCTCATCAC TTGACAATTA
4081 AGCCCCCATG CCTACCCTTG GTTTGGATGT CACACAGAGC TGAAGGGGTT GGGAGATGTC
4141 TCCCAACTGC TGATTAATGC TGAGTTTCTC AAAATTAACT TTTGTTCCTT TGGCTTAAAG
4201 CTCTGGCCCC AGCTGGGCCC TCCTGCTGCT CAGCAGTGCT TCCATGCATT CTCAATCAAC
4261 A**TTTGCTGG**C CCCTACCCTT TGGGTCTATT TATTGAGTGC TTACTATGTA CCAGGTACCA
4321 TGATGAGTGT TTCACACATT ATCTCATTTA ATCCTCTTGG TTCTACATTT AAGAATAGGT
4381 CCTAGTATCC TAATTTTACA CAGAAGGAGA AACCAAGGTT CAGAGAAACT AGAGCACTTG
4441 CCCAAGGTCT CACAGCTGAT ATACGGCTGG GCAAGAACCC TACCCTGGGG GGGCTCTCTA
4501 TCCCCAAAGT GTCTGCTTTC AAGCACCACA CCCACCCAGA TGACTTAGTC CATGGGAGAA
4561 ACGGAGGCCA CCAGGTGTGC AGCTCCCCAC CAATACTAAA AACATGTCCA CAGGCCAGGC
4621 TCTATGCTTC TCATTTAGGC CTCACAACAT TCCAGTG**AAG CAGAGGCTTA GA**AAAGTGAT
4681 CTCAGTTGGC CCAAGATGAC ATGAGTGGTA AATGAAGCCA CTGGAGTTCA AAGTCTGGCT
4741 CTCACGTCAG TTAAGCACAG GCCAGATTCC CTTTCTCTCC CTCCACATAG CTCTCACCTC
4801 CTCTGGCCTT TTTGCTGATG GCAGAGGTGT GCTCCTCTTT CTTCTTTCCA AGGGTACTTC
4861 CTCAACCCTG CTACAGTCCT TATCAACACA GCATTCATTC TCAGGGAATT GAAAGCTCCA
4921 AGTCACCCTT GTTACTGGCT CCTTCCATCA AGCAGACACC TCACCTAACC TGGAACATCC
4981 TTTCCACCAT GCTCTCCCCT TCATCAAGCT AACTACCTCT GCTCTTTTCC ATTGTTTCAA
5041 AACCTCTTGA AATGCAGCAG AAAGGAAAAG GAATAAACAT TTATTGAGAA CCTATTAGAC
5101 ACTTTACAGA TGTTATTTAA TTCTCACAAC CTTACCTATG AATTATCCCT GATTTACAGG

5161 TGAGCAAACA GGCTCAGAGA CACGGAACTT ACGGAGGTCT CAGAGCCAAT AAGCAACCAG
5221 GAGTAATCTG ACTCTAAAAC ATGTGCTGCC CCCATGCCAC CTCCATGACT CTACTCTCCC
5281 AAAGGTTGCT TGTGTCTACT AGCCAAATCC AAGGCCGCCT TCTTTGTCCT CACTCATCTG
5341 GTCCTCAGTG CCTCTACCAC TTACTTCTTC AACACTCCCC TCTCCA**GGTC ACA**CCCTCTC
5401 TGACCTTTTA TTCTCTTGCA TATGCATGTT CCTGAGAACA GCACTTAGTC CTCTTTCACT
5461 ACCCAATCTG TACTCCCTCG CAATCTCATC CTTCATCTCT CCTGATTTTC AAACATCCTA
5521 GACTGGATAT TTCAGACACT TAAAATTCAA CAGATCTCAA GAGAATATAT TTCTCATGAG
5581 CATCCTCCAT GGATCTCCCT GACCCGGATC TGTCTGCCTG TTCTAAGTCC AGGGCTCTTT
5641 GCATTATCCT ATGGGATGAA AGGCACCATC ATGGTCTGCC TGGCTGTGGA AAGCAAGAGG
5701 GACTCCTGGG CTCATCTGCC CTCCATAGGC AGGAACCCAA ATGGGGAAAG AGGAAGGGAA
5761 CTGTCTAAAA AAGTTGCTTT CTCAGGTTGT GTGGAGAGAC AGGACTCAGT GAGGATAGGC
5821 CCCTACAGGC CTATCAACCA GCCACTCTGT GGCTAAAAGT TAATTGTCCA ATAAAATTTA
5881 GCTATCCCAG GATAAGGCAT TTCTACTTTA TACGATTATT CTGTTTTAAA ATATATTTGC
5941 TTTGAGAGTA ATTCCCCAAA TAGAATTCCT ATTTTCTACA GCCTTGCTCT TCTTGTTTTA
6001 TCCTTATCAA TAGTGCCATC ATCCACTGCC CTGACACCTC TGATTCCTTC CTTTCCCTCC
6061 ATTTCACAAA ATCTATCAAT TCTACTTAGT GACTCTATCC CACTGGTGTT GAATTCTCCC
6121 TTTTTTTTTT AAACAGCAGA AACAGTTTAT TAAAATAGAA ATCTTATGTA GAACCTTTAT
6181 GCATAAAAAA GATTCAAGCT GCTTCACTTG AAAACAGGAA GCAGCACCAA ATCCCCCCAT
6241 ACTCAACT**CC AACCCTTGCT** TGTCTCAAGG CACCTACATA GAAACCCAGA GTGTCTCAGA
6301 AGAACAGCTG GAGCGCTACA GCCTGCTCTC TTCTGCTGTC CCCACTGCCC CCGGCTAGGA
6361 TCACCTCCTA CAAGATTCCT GCACAGCCTA CTGACTTTTC TCCCTGCTCA CCCTCTTCCA
6421 TCCCTCTAAC CCAGCTCCTA GCATCTAGTG CTTCTTGGAC TTTTGTACCA AGTAGCCCCA
6481 ACAGTAGGAG AGAGGTCTGG GGATAAGTGA AATGAATATG ACTATTTGAA TTACCACTAT
6541 CAGTTTTAGT ATCTTGAGGC TTCAAGAACT GCTCTACCTA GCACCACCAC CTTCCCTAAG
6601 GACATAAGTA CGTACTCTAG ACTGAAAGGT TCTACTATAC ACCATCTAAT CTGATCATTC
6661 CCCGGTTCCT CATTACTTAC GAGAGAAAAA TCCAAATTTC TTATTTTAGC ATTCAAGGCA
6721 CTTTACATGC AGGGCTCTAC TTCCAAGCTG GGGCATGTGG TTCCAGAGAA AGAGAGGATA
6781 CCTCAGGACA GTATGATACA TCTTACCTAA TGGTAAGTAA TTTAAAATAT TAAATTTATA
6841 TTAAATAGGT AATATGGTGT TAAAATGCAA CACCTACGTG AATTTGTAAG GCAAAACAGT
6901 ACTTCAAAGA AATGTATATG CAGTGAGACA TTTCTAGCTT TATTTCCAGA TCCCTTTCAA
6961 CCCAGGTACC ATTTAATCTC TGTAATTAGG TGTACTCTGG TGGAACACTC ACCATTTTTT
7021 AGATGAGGAA ACAGGGTCAG AAAGGTCAAG TGACCTATCC AA**GGTCACA**C AACTAGCAAG
7081 CAGTTAAGCT AGACTTTTAC TGTAGGCAGT CTGCCTCCAG TCCCGCTCCT AACCACTGTA
7141 CTGTATTATC TCCTCAAATG CATTAGAATC TTCACTTCCA TAAAGAAATA TGCTCTTTGT
7201 TTTTTTCTTT TGGAACACAA GGTTTAGTCC ATCACTCACT CACTTTTCCA CTTTTGATAA
7261 GATGTAAATA TTGAGAACTA TTTCTCTTCT CCAGTAGAAG **AAGAGCAACA GGCC**ATGTGT
7321 GATGGTAAAT GGCAGCTGAC ACTTGCGACA GTATCAAGAA GGCACTAAGG AATGGTGAGA
7381 AGGATGACTT GCCCTTTCTA AAGTGGCATC TCAGCTTGAG TCTGCAGAAT GCCTTGCAAA
7441 TCCAGCTCAG TACTGCTAGA TAATCCATAT TGACAAGAGA AGCTAAAGAT TCAGATTTTC
7501 TATGTGAAGT TTCCTGGTTT TAAAATGTTA GCAACAAATT CCAAATTTTT ACAGAACACT
7561 GTACAGGCCA GAAAGAACAA TCTGTGTGCT ACCAGTTTAC AATTAGCTAT GCTTCTTCCT
7621 CCTCAGACCT GGCA**TGCCCT TT**CCACCCCC TTGCCTTTGC TCTTAGTACC CCGTGTACAC
7681 TCCAACCTCT TCCACCTACA AATTTCTAAA TGCACAGTTC AAATGTTAGT TCTTCAGATT
7741 CCTGTCTCTG TAATAACACT TGCCCCTTTG TATAACAGTT CTCATGTATA TCTTCCTATT
7801 AGACCATGAG CTCCTGTGGT AGAAATTGTG TATTCATTAC TGTATCCCTA GTGTGCAGTA
7861 GATCACA GTAG

**k.** *PXDN* 3'UTR sequence with bioinformatically predicted hsa-miR-7973 target sequences. microT CDS v5.0 predicted target sequence locations are marked in green: positions 12-31 and 329-354. TargetScan and miRDB commonly predicted seed sequence location is marked in red: position 2298-2305.

1 GCTCCTGGGA G**GCTCCTCAG AGTTTGTCTG C**TGTGCCATC GTGAGATCGG GTGGCCGATG
61 GCAGGGAGCT GCGGACTGCA GACCAGGAAA CACCCAGAAC TCGTGACATT TCATGACAAC

121 GTCCAGCTGG TGCTGTTACA GAAGGCAGTG CAGGAGGCTT CCAACCAGAG CATCTGCGGA
181 GAAGGAGGCA CAGCAGGTGC CTGAAGGGAA GCAGGCAGGA GTCCTAGCTT CACGTTAGAC
241 TTCTCAGGTT TTTATTTAAT TCTTTTAAAA TGAAAAATTG GTGCTACTAT TAAATTGCAC
301 AGTTGAATCA TTTAGGCGCC TAAATTGAT**T TTGCCTCCCA ACACCATTTC TTTTT**AAATA
361 AAGCAGGATA CCTCTATATG TCAGCCTTGC CTTGTTCAGA TGCCAGGAGC CGGCAGACCT
421 GTCACCCGCA GGTGGGGTGA GTCTTGGAGC TGCCAGAGGG GCTCACCGAA ATCGGGGTTC
481 CATCACAAGC TATGTTTAAA AAGAAAATTG GTGTTTGGCA AACGGAACAG AACCTTTGAT
541 GAGAGCGTTC ACAGGGACAC TGTCTGGGGG TGCAGTGCAA GCCCCCGGCC TCTTCCCTGG
601 GAACCTCTGA ACTCCTCCTT CCTCTGGGCT CTCTGTAACA TTTCACCACA CGTCAGCATC
661 TAATCCCAAG ACAAACATTC CCGCTGCTCG AAGCAGCTGT ATAGCCTGTG ACTCTCCGTG
721 TGTCAGCTCC TTCCACACCT GATTAGAACA TTCATAAGCC ACATTTAGAA ACAGGTTTGC
781 TTTCAGCTGT CACTTGCACA CATACTGCCT AGTTGTGAAC CAAATGTGAA AAAACCTCCT
841 TCATCCCATT GTGTATCTGA TACCTGCCGA GGGCCAAGGG TGTGTGTTGA CAACGCCGCT
901 CCCAGCCGGC CCTGGTTGCG TCCACGTCCT GAACAAGAGC CGCTTCCGGA TGGCTCTTCC
961 CAAGGGAGGA GGAGCTCAAG TGTCGGGAAC TGTCTAACTT CAGGTTGTGT GAGTGCGTTA
1021 AAAAAAAAAA AAAAAAAGAA TCCCTATACC TCATTTGTAT TTTTAAAATG CGTGATGTTT
1081 TATGAAATTG TGTCCATTTT TTAGGTATTA GATATGGCAG AAAAACCATT TCCACTATGC
1141 AAAGTTCTTT TAGACGTCAG TGAAAATCAA CTCTCATACC TCATGGTCTC TCTTTAATTG
1201 ACCAAAACCT TCCATTTTTC TCTAAATACA AAGCGATCTG TGTTCTGAGC AACCTTTCCC
1261 CGAACACACA GCTTCAGTGC AGCACGCTGA CCTGAGTATC CACCATGTGC CAGGCACAGT
1321 GCTGGGCACA CGAGGCACCA AGGTCCGGGC CACCTGCCCG CAGCAAGGCC CAGCTGAGGT
1381 GGTGGAGGGA GCCCCTGAGG TCAGGGGCCG TTTCGGTTCA GGGTGGCAGG TGTCCAGCAC
1441 TGGGGTATGG CGTCGAGGCT TCCATGGGGT GGGGGAGGCC AGCTTCCTTC TGACAGGATG
1501 GGCGCATACA GTGCCTGGTG TGATTTGTGC ACAACCCGTG TTCCAGGTGC ACATCCTCCC
1561 AAGGAGACAC CCAGACCCTT CCAGCACGGG CCGGCCAAGT TGCTGCGGCG GAGGCAGCAT
1621 TTCAGCTGTG AGGAAGGTCA TTGGATTCAT GTGTTTTATC TGTAAAAATG GTTGTCTTAA
1681 CTTCTTAACC TCATATTGGT AAGTGATTGA TAAAAATTGG TTGGTGTTTC ATGACATGTG
1741 GACTTCTTTT GAAATAGCAA GTCAAATGTA GTGACCAAAT TGTGGAAGAG ATTTCTGTCA
1801 AATAGGAAAT GTGTAAGTTC GTCTAAAAGC TGATGGTTAT GTAAGTTGCT CAGGCACTCA
1861 GATGACAGCA GATTCTGGGT TCTGGGAGTG TTCTGTGCCT CTTACATGCC CTGGAGGCCT
1921 CATGGTCTCA GTGCTGAGGC GGCACACCTG TAGCACACCT GCGTAATGTG CGGTCTGGGC
1981 CAGTCACAAG GAATTGTGTT GTCTAAGCCA AAGGGGGAAG CTGACTGTGA TTTACCAAAA
2041 AAAATTCTGT AATTCAAACC AAAATGTCTG CGGAATCACC AGTTTGATAC TCTCTGTAAT
2101 CAGAACAGTG GGCAGTGCCT GGGTGAACGT GTCTAGCAGC CACTGTGCGG GATCGCTGTA
2161 ACAGGAGTGG AATGTACATA TTTATTTACT TTTCTAACTG CTCCAACAGC CAAATGCCTT
2221 TTTTATGACC ATTGTATTCA GTTCATTACC AAAGAAATGT TTGCACTTTG TAATGATGCC
2281 TTTCAGTTCA AATAAAT**GGG TCAC**ATTTTC AAATGGA

**l.** *TGFBR2* 3'UTR sequence with bioinformatically predicted target sequences for hsa-miR548ba and hsa-miR7973. TargetScan and miRDB commonly predicted target seed sequence location for hsa-miR-7973 at position 107-113 is marked in red. miRDB predicted seed sequence location for hsa-miR-548ba at position 810-816 is marked in yellow. TargetScan predicted seed sequence location for hsa-miR-548ba at position 1470-1476 is marked in blue.

1 CTCTTCTGGG GCAGGCTGGG CCATGTCCAA AGAGGCTGCC CCTCTCACCA AAGAACAGAG
61 GCAGCAGGAA GCTGCCCCTG AACTGATGCT TCCTGGAAAA CCAAGG**GGGT CAC**TCCCCTC
121 CCTGTAAGCT GTGGGGATAA GCAGAAACAA CAGCAGCAGG GAGTGGGTGA CATAGAGCAT
181 TCTATGCCTT TGACATTGTC ATAGGATAAG CTGTGTTAGC ACTTCCTCAG GAAATGAGAT
241 TGATTTTTAC AATAGCCAAT AACATTTGCA CTTTATTAAT GCCTGTATAT AAATATGAAT
301 AGCTATGTTT TATATATATA TATATATATC TATATATGTC TATAGCTCTA TATATATAGC
361 CATACCTTGA AAAGAGACAA GGAAAAACAT CAAATATTCC CAGGAAATTG GTTTTATTGG
421 AGAACTCCAG AACCAAGCAG AGAAGGAAGG GACCCATGAC AGCATTAGCA TTTGACAATC
481 ACACATGCAG TGGTTCTCTG ACTGTAAAAC AGTGAACTTT GCATGAGGAA AGAGGCTCCA
541 TGTCTCACAG CCAGCTATGA CCACATTGCA CTTGCTTTTG CAAAATAATC ATTCCCTGCC

601 TAGCACTTCT CTTCTGGCCA TGGAACTAAG TACAGTGGCA CTGTTTGAGG ACCAGTGTTC
661 CCGGGGTTCC TGTGTGCCCT TATTTCTCCT GGACTTTTCA TTTAAGCTCC AAGCCCCAAA
721 TCTGGGGGGC TAGTTTAGAA ACTCTCCCTC AACCTAGTTT AGAAACTCTA CCCCATCTTT
781 AATACCTTGA ATGTTTTGAA CCCCACTTT**T TACCTT**CATG GGTTGCAGAA AAATCAGAAC
841 AGATGTCCCC ATCCATGCGA TTGCCCCACC ATCTACTAAT GAAAAATTGT TCTTTTTTTC
901 ATCTTTCCCC TGCACTTATG TTACTATTCT CTGCTCCCAG CCTTCATCCT TTTCTAAAAA
961 GGAGCAAATT CTCACTCTAG GCTTTATCGT GTTTACTTTT TCATTACACT TGACTTGATT
1021 TTCTAGTTTT CTATACAAAC ACCAATGGGT TCCATCTTTC TGGGCTCCTG ATTGCTCAAG
1081 CACAGTTTGG CCTGATGAAG AGGATTTCAA CTACACAATA CTATCATTGT CAGGACTATG
1141 ACCTCAGGCA CTCTAAACAT ATGTTTTGTT TGGTCAGCAC AGCGTTTCAA AAAGTGAAGC
1201 CACTTTATAA ATATTTGGAG ATTTTGCAGG AAAATCTGGA TCCCCAGGTA AGGATAGCAG
1261 ATGGTTTTCA GTTATCTCCA GTCCACGTTC ACAAAATGTG AAGGTGTGGA GACACTTACA
1321 AAGCTGCCTC ACTTCTCACT GTAAACATTA GCTCTTTCCA CTGCCTACCT GGACCCCAGT
1381 CTAGGAATTA AATCTGCACC TAACCAAGGT CCCTTGTAAG AAATGTCCAT TCAAGCAGTC
1441 ATTCTCTGGG TATATAATAT GATTTTGAC**T ACCTTA**TCTG GTGTTAAGAT TTGAAGTTGG
1501 CCTTTTATTG GACTAAAGGG GAACTCCTTT AAGGGTCTCA GTTAGCCCAA GTTTCTTTTG
1561 CTTATATGTT AATAGTTTTA CCCTCTGCAT TGGAGAGAGG AGTGCTTTAC TCCAAGAAGC
1621 TTTCCTCATG GTTACCGTTC TCTCCATCAT GCCAGCCTTC TCAACCTTTG CAGAAATTAC
1681 TAGAGAGGAT TTGAATGTGG GACACAAAGG TCCCATTTGC AGTTAGAAAA TTTGTGTCCA
1741 CAAGGACAAG AACAAAGTAT GAGCTTTAAA ACTCCATAGG AAACTTGTTA ATCAACAAAG
1801 AAGTGTTAAT GCTGCAAGTA ATCTCTTTTT TAAAACTTTT TGAAGCTACT TATTTTCAGC
1861 CAAATAGGAA TATTAGAGAG GGACTGGTAG TGAGAATATC AGCTCTGTTT GGATGGTGGA
1921 AGGTCTCATT TTATTGAGAT TTTTAAGATA CATGCAAAGG TTTGGAAATA GAACCTCTAG
1981 GCACCCTCCT CAGTGTGGGT GGGCTGAGAG TTAAAGACAG TGTGGCTGCA GTAGCATAGA
2041 GGCGCCTAGA AATTCCACTT GCACCGTAGG GCATGCTGAT ACCATCCCAA TAGCTGTTGC
2101 CCATTGACCT CTAGTGGTGA GTTTCTAGAA TACTGGTCCA TTCATGAGAT ATTCAAGATT
2161 CAAGAGTATT CTCACTTCTG GGTTATCAGC ATAAACTGGA ATGTAGTGTC AGAGGATACT
2221 GTGGCTTGTT TTGTTTATGT TTTTTTTTCT TATTCAAGAA AAAAGACCAA GGAATAACAT
2281 TCTGTAGTTC CTAAAAATAC TGACTTTTTT CACTACTATA CATAAAGGGA AAGTTTTATT
2341 CTTTTATGGA ACACTTCAGC TGTACTCATG TATTAAAATA GGAATGTGAA TGCTATATAC
2401 TCTTTTTATA TCAAAAGTCT CAAGCACTTA TTTTTATTCT ATGCATTGTT TGTCTTTTAC
2461 ATAAATAAAA TGTTTATTAG ATTGAATAAA GCAAAATACT CAGGTGAGCA TCCTGCCTCC
2521 TGTTCCCATT CCTAGTAGCT AAA
